# *Plat* safeguards maternally aged oocytes against programmed cell death through activating the Erk1/2 pathway

**DOI:** 10.1101/2025.02.11.637629

**Authors:** Xingsi He, Hanwen Zhang, Ya Wang, Huanyu Yan, Qiuzhen Chen, Min Su, Qiaozhen Shi, Xiao Zeng, Wei Sheng, Yangmin Wang, Chikun Wang, Shuyue Hou, Zhibin Hu, Yuanlin He, Xi Wang

## Abstract

The decline in oocytes quality and developmental potential with female reproductive aging is well recognized, yet the underlying mechanisms remain insufficiently investigated. In this study, through an integrative analysis of transcriptomes and morphologies of individual oocytes from young and aged mice, morphologically defective aged oocytes are identified with distinct transcriptomic features. Further analysis demonstrates that both apoptotic and ferroptotic pathways are activated in the defective aged oocytes, and simultaneously blocking both pathways reverses the defective morphology to the largest extent. The *Plat* gene, which encodes tissue-type plasminogen activator (tPA), is downregulated with oocyte aging, and *Plat* knockdown increases oocytes susceptibility to both apoptosis and ferroptosis. Mechanistically, tPA functions as an upstream signaling molecule for Erk1/2 activation by interacting with particular phosphorylation kinases such as Alk. Consequently, *Plat* loss downregulates Erk1/2 pathway activity in oocytes, leading to degeneration through programmed cell death. Supplementing exogenous tPA in *in vitro* oocyte maturation cultures reduces defect rate of aged oocytes, thereby improving oocyte quality and developmental potential. Collectively, *Plat* plays a pivotal role in protecting aged mouse oocytes from programmed cell death, and tPA supplementation may serve as a potential clinical strategy to enhance oocyte quality in females of advanced maternal age.

## INTRODUCTION

Deterioration of oocyte quality is a hallmark of reproductive aging and a primary factor contributing to the fertility decline in women of advanced maternal age ^1^, a trend that has become increasingly prevalent due to socioeconomic shifts toward later childbearing ^2,3^. Consequently, disentangling the molecular mechanisms that link aging with the degradation of oocyte quality, and developing targeted interventions to mitigate these effects, has become a pressing research priority for preserving the reproductive potential in aging women.

Morphological abnormalities in oocytes, such as fragmentation and cytoplasmic darkening, are positively correlated with maternal age^4,5^, and are observed across multiple mammalian species, including human and mouse^6,7^. These abnormal features often serve as early indicators of adverse pregnancy outcomes^8–10^, and may reflect disruptions in subcellular and molecular homeostasis within the oocyte. Specifically, dysregulation in processes such as mitophagy^11^, cohesin maintenance^12^, and oxidative stress response^13,14^ have been associated with these morphological changes. While oocyte degeneration, particularly fragmentation, has traditionally been attributed to apoptotic cell death triggered by DNA damage, metabolic imbalance, and other signaling disruptions^15,16^, recent studies suggest that additional types of programmed cell death (PCD), such as ferroptosis, may also play a role. Iron accumulation in aged oocytes, for instance, has been shown to damage the redox homeostasis, compromising oocyte quality^17^. Very recently, Zhang. et al have demonstrated that oocyte ferroptosis can be activated in the absence of certain protective genes^18^, underscoring that oocyte degeneration may involve multiple PCD pathways beyond apoptosis. However, detailed studies on the PCD mechanisms associated with oocyte aging and the factors that trigger such pathways remain largely unresolved.

In this study, we employ single-oocytes transcriptome sequencing to investigate molecular changes in aged oocytes compared to those from young mice. Our transcriptomic analysis links gene expression alterations with morphological abnormalities, and in-depth examination of differentially expressed genes indicates that defective oocytes are undergoing PCD, involving both apoptotic and ferroptotic pathways. We identify tissue-type plasminogen activator as a prominently down-regulated gene in defective oocytes, with evidence suggesting it acts upstream of the Erk1/2 pathway, a critical pathway involved in oocyte meiotic maturation^19^. Supplementation of this protein appears to ameliorate oocyte degeneration, pointing to potential therapeutic strategies to counteract female reproductive aging.

## RESULTS

### Single-oocyte RNA-seq identifies a subgroup of aged oocytes exhibiting morphological abnormalities

It has been reported that maternal fertility declines with reproductive aging, primarily due to reduced oocyte quantity and quality^20^. We confirmed this phenomenon by comparing young (3-8 weeks) and aged mice (10-12 months), examining the number of ovulated oocytes (Supplementary Fig. S1A), fertilization rate (Supplementary Fig. S1B), and blastocyst formation rate (Supplementary Fig. S1C) following in vitro fertilization (IVF) experiments.

To investigate the molecular changes underlying oocyte aging, we collected metaphase II (MII) oocytes from both young and aged mice for both morphological analysis and single-oocyte RNA sequencing (soRNA-seq) (Fig. 1A). Since soRNA-seq was able to profile transcriptomes of individual oocytes, here we took all the oocytes ovulated from young and aged mice for analysis, including those exhibiting morphological defects. Bioinformatics analysis of the soRNA-seq data revealed distinct transcriptomic differences between the young and aged oocytes. Using unsupervised clustering, we identified three main clusters, two of which, C1 and C2, were predominantly composed of aged oocytes (Fig. 1B). Further morphological examining revealed that one of the aged clusters, C2, contained exclusively fragmented and darkened oocytes, whereas the other cluster, C1, comprised oocytes with intact morphology (Fig. 1C). While the majority of oocytes in C2 displayed fragmentation, a more advanced state of cellular degradation, darkened oocytes were likely at a pre-fragmentation stage, given their similarity in transcriptomic profiles. Notably, these morphological defects were observed only in aged oocytes, suggesting they were characteristics of age-related oocyte degeneration. Based on the connection between expression profiles and structural morphology, we therefore designated the three clusters of oocytes as Young (C0), Aged-Normal (C1) and Aged-Defective (C2).

**Fig. 1.**
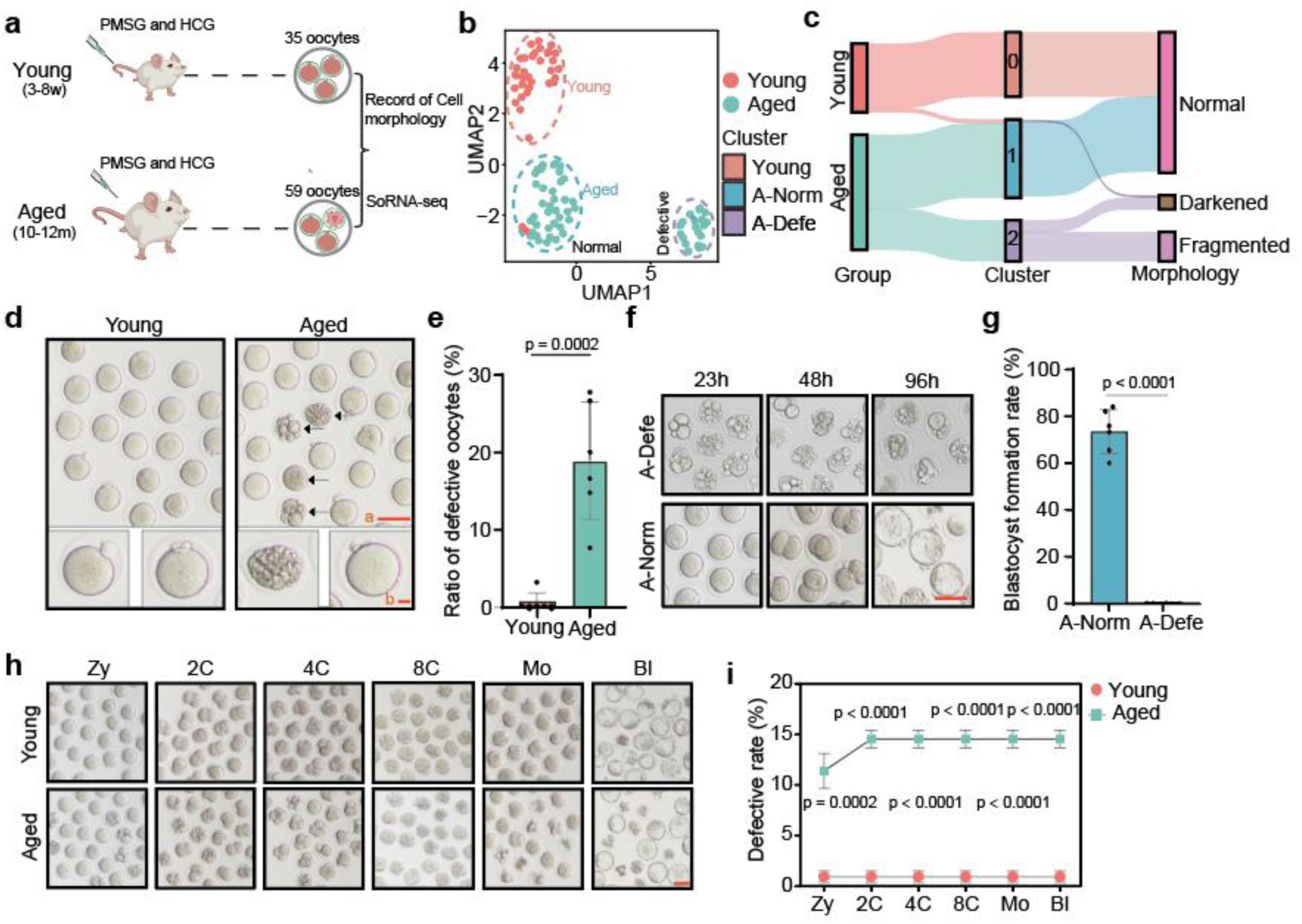
A fraction of aged oocytes displayed morphological defects. (**A**) A total of 94 oocytes extracted from young (3-8 wk) and aged mice (10-12 m), subject to morphological recording and single-oocyte RNA-seq (soRNA-seq) profiling. (**B**) Uniform manifold approximation and projection (UMAP) plot showing oocyte clusters. Points depict oocytes, and the dashed circles represent clusters. Colors of the points represent oocytes from young or aged mice. (**C**) Sankey plot delineating correspondence across mouse age groups, oocyte clusters, and cellular morphologies. (**D**) Representative images showing extruded oocytes from young and aged mice. Arrows: defective oocytes. Scale bars, a: 100μm, b: 20μm. (**E**) Barplots showing the defect rates of oocytes from young and aged mice. The data are represented as the mean ± s.d. from 6 unique biological samples. *** P < 0.001 according to two-tailed unpaired t-test. (**F**) Representative images showing the developmental stages of embryos derived from morphologically normal (A-Norm) and defective (A-Defe) oocytes of aged mice. Stages at different time points: 23h -Zygote, 48h - 2-cell, 96h - blastocyst. (**G**) Barplots comparing the blastocyst formation rate developed from morphologically normal and defective MII-stage oocytes, as exemplified in (F). Data are demonstrated as mean ± s.d. of 6 biologically independent samples. **** P < 0.0001 according to two-tailed unpaired t-test. (**H**) Representative images of embryo development at various stages following IVF from oocytes of young and aged mice with the morphologically defective oocytes of aged mice excluded. Scale bar: 100 μm. (**I**) Cumulative defect rates at each developmental stage from young and aged oocytes, as shown in (H). The data are displayed as the mean ± s.e.m. from 6 biological replicates. *** P < 0.001 and **** P < 0.0001 according to two-tailed unpaired t-test. ZY, zygote; 2C, 2-cell; 4C, 4-cell; 8C, 8-cell; Mo, morula; BL, blastocyst.

On the morphological side, freshly ovulated oocytes from aged mice exhibited a high incidence of morphological defects, with oocyte fragmentation being notably prevalent (Fig. 1D). The defective ratio in aged oocytes reached up to 20% on average, whereas such morphological defects were rarely observed in oocytes from young mice (Fig. 1E). Defective aged oocytes displayed notable cytoplasmic granularity and nuclear debris (Supplementary Fig. S1D). In assessing oocyte developmental capacity, we observed that defective oocytes had no potential for embryonic progression through IVF (Fig. 1F), as confirmed across replicated experiments (Fig. 1G). Conversely, aged oocytes of normal morphology at the MII stage were capable of fertilization and could progress to the blastocyst stage (Fig. 1H), with only a minor subset showing defects at the zygote (∼11.5%) or 2-cell (∼3.3%) stages (Fig. 1I). Given that morphological abnormalities of embryos developed from normal aged oocytes primarily emerged at the MII and zygote stages, we hypothesized that these defects were likely linked to maternal factors, further underscoring our focus on MII oocytes in this study. These analyses also highlight that the declined fertilization rate (Supplementary Fig. S1B) and blastocyst formation rate (Supplementary Fig. S1C) in aged oocytes were primarily attributed to the incidence of defective oocytes. Therefore, lowering the defect rates of aged oocytes could improve the fertility for aged females.

### Aged oocytes exhibit widespread transcriptomic alterations

Oocytes in the Aged-Defective cluster exhibited a substantial number of differentially expressed genes (DEGs) when compared to Young oocytes (Supplementary Fig. S2A) and Aged-Normal oocytes (Supplementary Fig. S2B). As expected, the number of DEGs detected between Aged-Normal and Young oocytes was relatively smaller, but their divergence was still obvious (Supplementary Fig. S2C). The up- and down-regulated genes in Aged-Defective oocytes, compared to either Young or Aged-Normal groups, exhibited significant overlap (Supplementary Fig. S2D,E), indicating that transcriptomic alterations in the Aged-Defective oocytes were more specific. Notably, the number of downregulated genes was more than twice that of upregulated genes in Aged-Defective oocytes, suggesting a possible global reduction in gene expression during oocyte degeneration (Supplementary Fig. S2A,B). This hypothesis was further supported by a significant decrease in the number of genes with detectable RNA expression in Aged-Defective oocytes in comparison to the other groups (Fig. 2A).

**Fig. 2.**
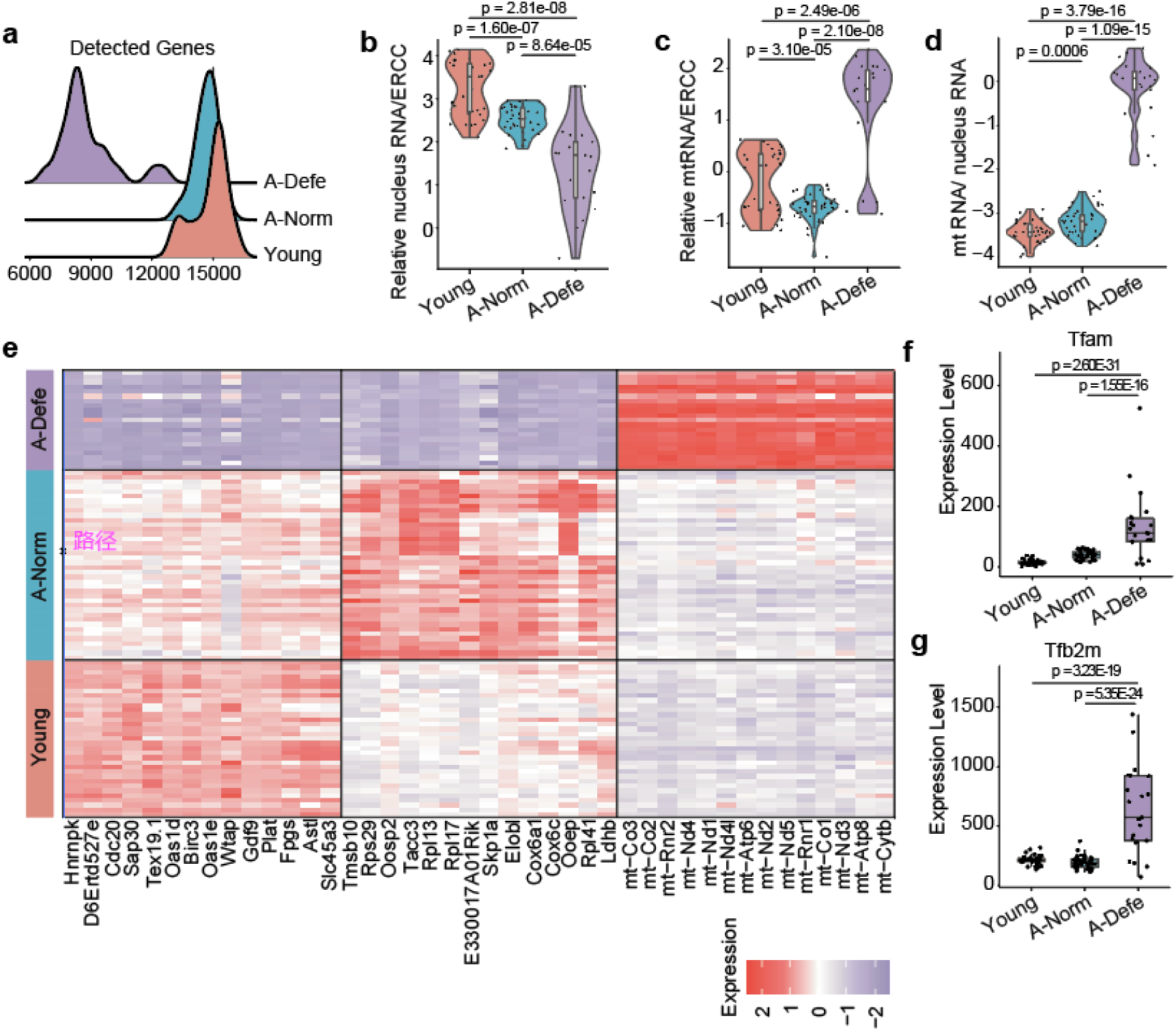
soRNA-seq data reveal gene expression alterations between oocyte clusters. (**A**) Density plots illustrating the distribution of total numbers of expressed genes in different oocyte clusters. (**B**) Violin plots displaying nuclear RNA expression relative to ERCC RNA of the three oocyte clusters. Expression levels of nuclear RNA and ERCC RNA were quantified in FPKM, and log-transformed. **** P < 0.0001 according to two-tailed unpaired t-test. (**C**) Violin plot comparing mitochondrial RNA expression relative to ERCC RNA across the three oocyte clusters. Expression levels of mitochondrial RNA and nuclear RNA were quantified in FPKM, and log-transformed. **** P < 0.0001 according to two-tailed unpaired t-test. (**D**) Violin plot comparing the ratio of mitochondrial RNA expression to nuclear RNA expression in the three oocyte clusters. Expression levels of mitochondrial RNA and ERCC RNA were quantified in FPKM, and log-transformed. *** P < 0.001 and **** P < 0.0001 according to two-tailed unpaired t-test. (**E**) Heatmap showing the marker gene expression across the three oocyte clusters. In each cluster, the top 14 marker genes are shown, with respectively gene names indicated. (**F**) Boxplots comparing *Tfam* expression across the three oocyte clusters. **** P < 0.0001 according to the Wald test implemented in DESeq2. (**G**) Boxplots comparing *Tfb2m* expression across the three oocyte clusters. **** P < 0.0001 according to the Wald test implemented in DESeq2.

To explore the expression alteration further, we partitioned RNA transcripts into nuclear and mitochondrial fractions based on their genomic origins, and employed External RNA Control Consortium (ERCC) spike-ins to normalize RNA molecule quantification across oocytes. Interestingly, this analysis revealed a reduction in nuclear RNA expression (Fig. 2B; Supplementary Fig. S2F) alongside an increase in mitochondrial RNA (mtRNA) expression in Aged-Defective oocytes (Fig. 2C; Supplementary Fig. S2G). Given the predominance of nuclear-transcribed genes, this finding aligns with the observed global decrease in overall gene expression in Aged-Defective oocytes. While Aged-Normal oocytes showed decreased expression in nuclear RNA, they also displayed a reduction in mtRNA expression, with the latter on the contrary of Aged-Defective oocytes. Notably, the ratio of mtRNA expression to nuclear RNA expression was able to better separate normal and defective oocytes, and Aged-Normal oocytes lay in between young and defective oocytes (Fig. 2D). The significant increase in mtRNA expression in Aged-Defective oocytes suggests a potential link to metabolic dysregulation induced by cellular stress and may be associated with specific programmed cell death (PCD) pathways.

Cluster-specific marker gene analysis revealed that *Gdf9* and *Astl* were specifically high expressed in Young oocytes, translation-related genes such as *Rps29* and *Rpl17* were elevated to some extent in Aged-Normal oocytes, and mitochondrial gene expression was substantially elevated in Aged-Defective oocytes (Fig. 2E). The elevated mitochondrial genes included those critical for the NADH dehydrogenase complex (e.g. *mt-Nd1*, *mt-ND4*), complex IV in the mitochondrial respiratory chain (*Co1* and *Co2*), and complex V subunits (*mt-Atp6* and *mt-Atp8*)^21^. Furthermore, *Tfam* and *Tfb2m*, two key transcription factors governing mitochondrial transcription^22^, were markedly upregulated in Aged-Defective oocytes (Fig. 2F,G), suggesting that the elevation of mtRNA expression was likely driven by enhanced transcriptional activity of mitochondrial genes. Gene ontology (GO) enrichment analysis of DEGs between oocyte clusters further highlighted upregulated apoptotic processes and disruptions in lipid metabolism within Aged-Defective oocytes (Supplementary Fig. S2H-K). In summary, our soRNA-seq analysis revealed a significant increase in mtRNA transcription in defective oocytes, potentially linking these changes to mitochondria-dependent PCD pathways.

### Apoptosis contributes to morphological defects in aged oocytes

To verify whether the apoptotic pathway was activated in defective oocytes, we first examined the expression of key apoptosis-related genes using the soRNA-seq data. Pro-apoptotic genes such as *Trp53inp1* and *Tnfaip8*^23,24^ were significantly upregulated in Aged-Defective oocytes, while anti-apoptotic genes, including *Bcl2l10* and *Bcl2l12*^25,26^, were markedly downregulated (Fig. 3A). To further validate these findings, we performed a terminal deoxynucleotidyl transferase dUTP nick end labeling (TUNEL) assay to assess DNA fragmentation levels, a hallmark of apoptosis^27^. TUNEL fluorescence intensity increased substantially in aged oocytes, with the highest levels observed in the defective subgroup (Fig. 3B,C). We next employed Annexin V/PI dual staining to determine the apoptotic stage across oocyte groups. In early apoptotic cells, externalized phosphatidylserine is detectable by Annexin V conjugated to red fluorophores, while intact plasma membranes prevent the entry of propidium iodide (PI). In late apoptosis, compromised membranes permit PI entry, resulting in green nuclear fluorescence. This staining revealed that aged oocytes with normal morphology were primarily in the early apoptotic stage, whereas defective oocytes had progressed to late-stage apoptosis (Fig. 3D,E).

**Fig. 3.**
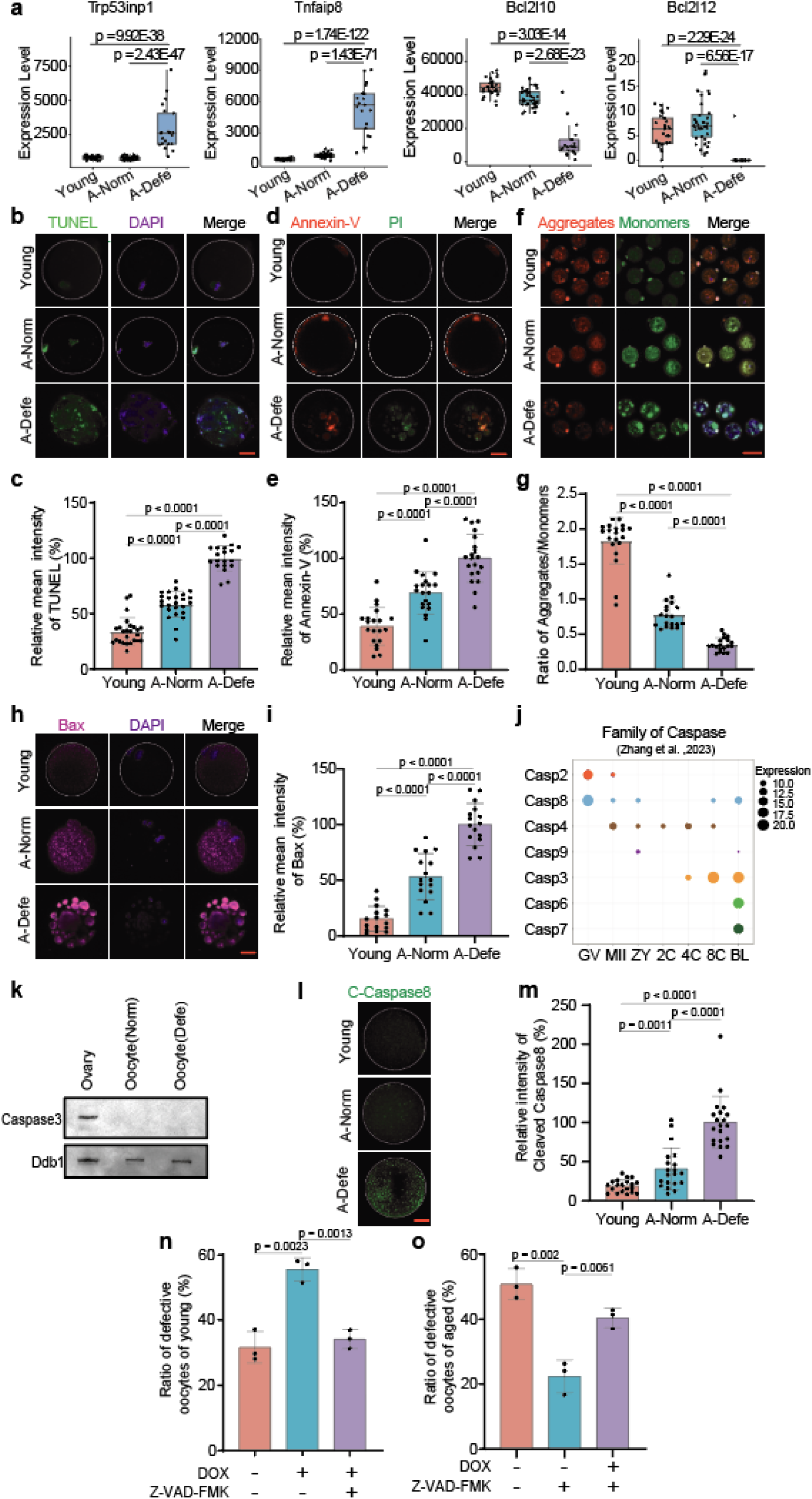
Characterization of apoptosis in aged oocytes. (**A**) Boxplots comparing the expression levels of representative apoptotic genes in Young, A-Norm and A-Defe oocytes. **** P < 0.0001 according to the Wald test implemented in DESeq2. (**B**) Representative images showing TUNEL levels in Young, A-Norm and A-Defe oocytes. Scale bar, 20 μm. (**C**) Barplots comparing relative fluorescence intensity of TUNEL in Young (n = 25), A-Norm (n = 25), and A-Defe (n = 19) oocytes. The data are represented as the mean ± s.d. **** P < 0.0001 according to two-tailed unpaired t-test. (**D**) Representative images showing Annexin-V and PI co-staining in Young A-Norm and A-Defe oocytes. Scale bar, 20 μm. (**E**) Barplots comparing relative fluorescence intensity of Annexin-V in Young (n = 20), A-Norm (n = 20), and A-Defe (n = 20) oocytes. The data are represented as the mean ± s.d. **** P < 0.0001 according to two-tailed unpaired t-test. (**F**) Representative images showing the assessment of mitochondrial membrane potential (ΔΨm) levels in Young, A-Norm, A-Defe oocytes using JC-1 staining (Red: high ΔΨm; Green: low ΔΨm). Scale bar, 100 μm. (**G**) Barplots comparing ratios of red to green fluorescence intensities in Young (n = 20), A-Norm (n = 19), and A-Defe (n = 19) oocytes. The data are represented as the mean ± s.d. **** P < 0.0001 according to two-tailed unpaired t-test. (**H**) Representative images showing Bax protein levels and distribution in Young, A-Norm, and A-Defe oocytes. Scale bar, 20 μm. (**I**) Barplots comparing relative fluorescence intensity of Bax in Young (n = 16), A-Norm (n = 16), and A-Defe (n = 16) oocytes. The data are represented as the mean ± s.d. **** P < 0.0001 according to two-tailed unpaired t-test. (**J**) Bubble chart depicting the relative expression patterns of caspase family proteins during oocyte maturation and early embryonic development. Protein expression quantified using log2-transformed intensity-based absolute quantification (iBAQ) values. ZY, zygote; 2C, 2-cell; 4C, 4-cell; 8C, 8-cell; BL, blastocyst. (**K**) Immunoblots showing Caspase3 expression in ovaries and oocytes (normal and defective). (**L**) Representative images showing Cleaved-Caspase8 expression levels and distribution in Young, A-Norm, and A-Defe oocytes. Scale bar, 20 μm. (**M**) Barplots comparing relative fluorescence intensity of Cleaved-Caspase8 in Young (n = 20), A-Norm (n = 20), and A-Defe (n = 20) oocytes. The data are represented as the mean ± s.d. ** P < 0.01 and **** P < 0.0001 according to two-tailed unpaired t-test. (**N**) Barplots comparing defect rates of oocytes from young mice treated with DMSO (n = 104), DOX only (n = 101), and DOX with Z-VAD-FMK (n = 94) for 48 hours. Data are represented as the average percentage of at least 3 independent experiments. The data are represented as the mean ± s.d. ** P < 0.01 according to two-tailed unpaired t-test. (**O**) Barplots comparing defect rates of oocytes from aged mice treated with DMSO (n = 85), Z-VAD-FMK only (n = 66), and DOX with Z-VAD-FMK (n = 70) for 48 hours. Data are represented as the average percentage of at least 3 independent experiments. The data are represented as the mean ± s.d. ** P < 0.01 according to two-tailed unpaired t-test.

Given the observed dysregulation of mtRNA expression in Aged-Defective oocytes, we assessed mitochondrial membrane potential using JC-1 staining across different oocyte groups. JC-1 forms red-fluorescent J-aggregates in high mitochondrial membrane potential conditions, but remains in its monomeric form and emits green fluorescence under low membrane potential^28^. The mitochondrial membrane potential index, calculated as the red / green fluorescence intensity ratio, showed a significantly reduction in aged oocytes, with defective oocytes exhibiting the most pronounced decline (Fig. 3F,G), indicating severe mitochondrial dysfunction. Given that mitochondrial dysfunction is known to trigger apoptosis through multiple mechanisms^29^, these findings suggest a link between mitochondrial impairment and apoptosis in defective aged oocytes. Consistently, Bcl-2-associated X protein (*Bax*), a key regulator of mitochondrial outer membrane permeabilization during apoptosis^30^, was significantly upregulated in aged oocytes, with the highest levels observed in defective oocytes (Fig. 3H,I).

The caspase family plays a pivotal role in the regulation of apoptosis^31^. Analysis of previously published proteomic data^32^ revealed that Caspase-3, an executioner caspase in the apoptotic pathway, was nearly undetectable in both GV and MII oocytes as well as in zygotes and 2-cell embryos, with its expression first appearing at the 4-cell stage (Fig. 3J). Western blot analysis further confirmed Caspase-3 protein expression in ovarian tissue but not in oocytes (Fig. 3K). Nevertheless, Caspase-8, a potential upstream mediator of apoptosis, was actively expressed at the GV, MII and zygote stages. Therefore, we hypothesized that apoptosis in defective aged oocytes was primarily mediated by Caspase-8 rather than Caspase-3. This hypothesis was additionally supported by Caspase-8 staining experiments, which demonstrated that cleaved Caspase-8 was significantly elevated in aged oocytes, reaching its maximum values in Aged-Defective oocytes (Fig. 3L,M).

To further explore the relationship between apoptosis with the morphological defects of oocytes, we treated young oocytes with doxorubicin (Dox), a topoisomerase inhibitor and potent apoptosis inducer. Dox treatment for 48 hours significantly increased the ratio of oocyte with defective morphology, and the ratio was markedly reduced by co-treatment with Z-VAD-FMK, a general caspase inhibitor^33^ (Fig. 3N; Supplementary Fig. S3A,B). Since aged oocytes were more prone to morphological defects, they likely exhibited greater susceptibility to apoptosis. Z-VAD-FMK treatment in aged oocytes for 48 hours drastically reduced morphological defect rates (Fig. 3O). Collectively, these findings clearly demonstrate that oocyte apoptosis contributes to morphological fragmentation in aged oocytes, and blocking the apoptotic pathway can effectively mitigate apoptotic cell death in aged oocytes.

### Defective aged oocytes also undergo ferroptosis

Revisiting our soRNA-seq data, the GO enrichment analysis revealed significant enrichment of lipid metabolic and catabolic processes among the downregulated genes in Aged-Defective oocytes (Supplementary Fig. S2J), pointing to the activation of ferroptosis, an additional form of PCD. Ferroptosis, an iron-dependent lipid peroxidation pathway, is closely tied to disruptions in lipid metabolism^34^. To investigate this point further, we applied COMPASS^35^ algorithm to predict metabolic flux based on our soRNA-seq data. While Young and Aged-Normal oocytes exhibited extensive metabolic activity divergence, comparing Aged-Defective oocytes to other groups showed indeed larger perturbations in lipid and amino acid metabolism (Supplementary Fig. S3C,D). In particular, a substantially increased activity of fatty acid synthesis and glutamate metabolism was evidenced in Aged-Defective oocytes, providing important metabolic basis for ferroptosis^36^ (Fig. 4A). Expression analysis of ferroptosis marker genes confirmed significant downregulation of *Gpx4* and *Hspb1*, as well as significant upregulation of *Ireb2* in Aged-Defective oocytes (Fig. 4B). Further exploration using FerrDb, the database of ferroptosis regulators^37^, demonstrated a general upregulation of ferroptosis drivers alongside a marked downregulation of ferroptosis suppressors in this oocyte subgroup (Fig. 4C; Supplementary Fig. S3E). The downregulation of *Gpx4* in defective oocytes was further validated based on immunofluorescence at the protein level (Fig. 4D,E).

**Fig. 4.**
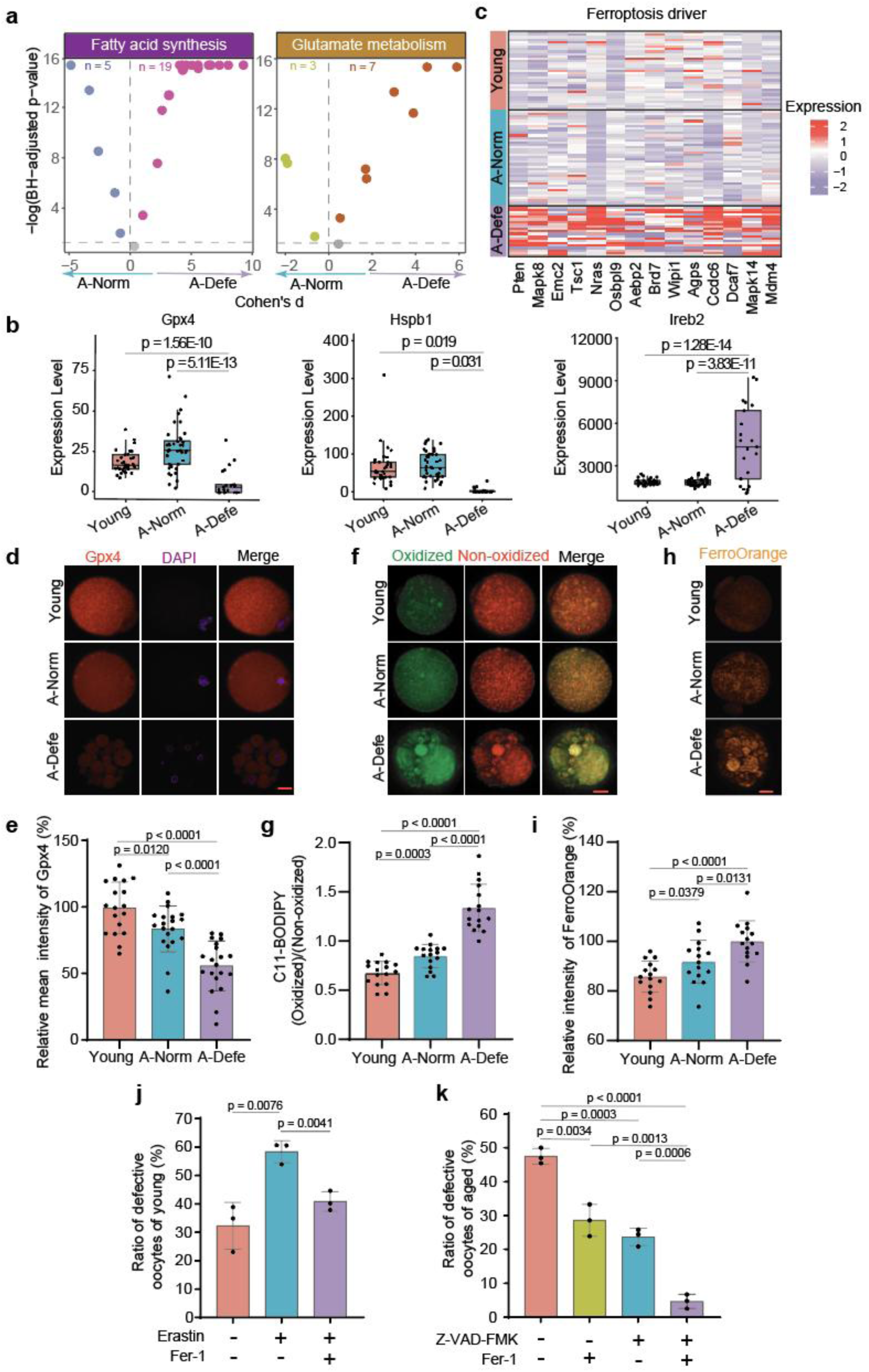
Characterization of ferroptosis in aged oocytes. (**A**) Volcano plot showing dissimilarity of COMPASS scores of individual reactions between A-Defe and A-Norm oocytes. Shown are two metabolic subsystems: Fatty acid synthesis (left) and glutamate metabolism (right). (**B**) Boxplots comparing the expression levels of representative ferroptosis marker genes (Gpx4, Hspb1, and Ireb2) in Young, A-Norm, and A-Defe oocytes. * P < 0.05 and **** P < 0.0001 according to the Wald test implemented in DESeq2. (**C**) Heatmap displaying the expression levels of selected ferroptosis driver genes. Oocytes were grouped according to the Young, A-Norm, and A-Defe clusters. (**D**) Representative images showing expression levels and distribution of Gpx4 proteins in Young, A-Norm, and A-Defe oocytes. Scale bar, 20 μm. (**E**) Barplots comparing relative fluorescence intensity of Gpx4 in Young (n = 20), A-Norm (n = 20), and A-Defe (n = 20) oocytes. The data are represented as the mean ± s.d. * P < 0.05 and **** P < 0.0001 according to two-tailed unpaired t-test. (**F**) Representative images showing the assessment of lipid peroxidation levels in Young, A-Norm, and A-Defe oocytes using C11-BODIPY staining (green: oxidized; red: non-oxidized). Scale bar, 20 μm. (**G**) Barplots comparing green / red fluorescence intensity ratios between Young (n = 16), A-Norm (n = 16), and A-Defe (n = 16) oocytes. The data are represented as the mean ± s.d. *** P < 0.001 and **** P < 0.0001 according to two-tailed unpaired t-test. (**H**) Representative images showing FerroOrange staining in Young, A-Norm, and A-Defe oocytes. Scale bar, 20 μm. (**I**) Barplots comparing relative fluorescence intensity of FerroOrange in oocytes of Young (n = 15), A-Norm (n = 15), and A-Defe oocytes (n = 15). The data are represented as the mean ± s.d. * P < 0.05 and **** P < 0.0001 according to two-tailed unpaired t-test. (**J**) Barplots comparing defect rates of young oocytes treated with DMSO (n = 85), Erastin only (n = 105), and Erastin with Fer-1 (n = 129) for 48 hours. The data are represented as the mean ± s.d. ** P < 0.01 according to two-tailed unpaired t-test. (**K**) Barplots comparing defect rates of aged oocytes treated with DMSO (n = 110), Z-VAD-FMK (n = 100), Fer-1 (n = 104) and a combination of Z-VAD-FMK and Fer-1 (n = 91) for 48 hours. The data are represented as the mean ± s.d. ** P < 0.01, *** P < 0.001 and **** P < 0.0001 according to two-tailed unpaired t-test.

To validate these findings functionally, we conducted lipid peroxidation assay in different groups of oocytes, which revealed a substantial elevation in lipid peroxidation levels in Aged-Defective oocytes, as well as a significantly yet smaller increase in Aged-Normal oocytes (Fig. 4F,G). Similarly, intracellular Fe^2+^ levels, assessed using FerroOrange staining, were significantly increased in aged oocytes, with the greatest increase observed in defective oocytes (Fig. 4H,I). To confirm the role of ferroptosis, we treated young oocytes with the ferroptosis agonist Erastin, which induced morphological defects. Co-treatment with the ferroptosis inhibitor Fer-1 significantly mitigated these effects, underscoring the contribution of ferroptosis to oocyte degeneration (Fig. 4J).

Finally, given that defective oocytes were likely undergoing both apoptotic and ferroptotic cell death, we evaluated aged oocyte survival under dual inhibition of apoptosis and ferroptosis by treating Z-VAD-FMK and Fer-1, respectively. While Z-VAD-FMK or Fer-1 alone reduced oocyte defects in aged oocytes, more than 20∼30% of oocytes remained defective (Fig. 4K). However, the combination of Z-VAD-FMK and Fer-1 dramatically decreased defect rates to below 5% (Fig. 4K). Collectively, the above results demonstrate that ferroptosis, alongside apoptosis, is a major contributor to morphological defects in aged oocyte, so that dual inhibition of these pathways markedly improves oocyte survival and reduces mortality.

### Programed cell death of aged oocytes attributed to loss of *Plat* expression

To elucidate the mechanisms underlying PCD in aged oocytes, particularly the defective ones, we re-examined the DEGs between the three clusters revealed by so-RNAseq (Fig. 1B). Among the downregulated genes in defective oocytes, *Plat* stood out as one of the most significantly decreased (Supplementary Fig. S4A,B); *Plat* was also downregulated in Aged-Normal oocytes when compared to Young cluster (FC=0.65, adjusted P = 2.55e-9; Fig. 5A). Furthermore, reanalysis of published proteomic data^30,38^, confirmed the tissue-type plasminogen activator (tPA) protein, encoded by *Plat*, was substantially upregulated in MII-stage oocytes compared to GV-stage oocytes, likely through translational activation, highlighting its important functions in MII-stage oocytes (Fig. 5B; Supplementary Fig. S4C,D). Consistently, our immunofluorescence assays validated a sharp increase in tPA protein levels at the MII stage (Supplementary Fig. S4E). When considering genes satisfactory with both downregulation in aged oocytes and protein-level upregulation at the MII stage, we found 21 genes with *Plat* ranked at the top (Fig. 5C). The above results suggest that *Plat* is a critical maternal factor playing critical roles in MII oocytes. Notably, immunofluorescence assays confirmed that tPA protein level was downregulated in aged oocytes, with markedly reduced in defective aged oocytes (Fig. 5D,E), implicating *Plat* loss was likely contributed to the defects observed in aged oocytes.

**Fig. 5.**
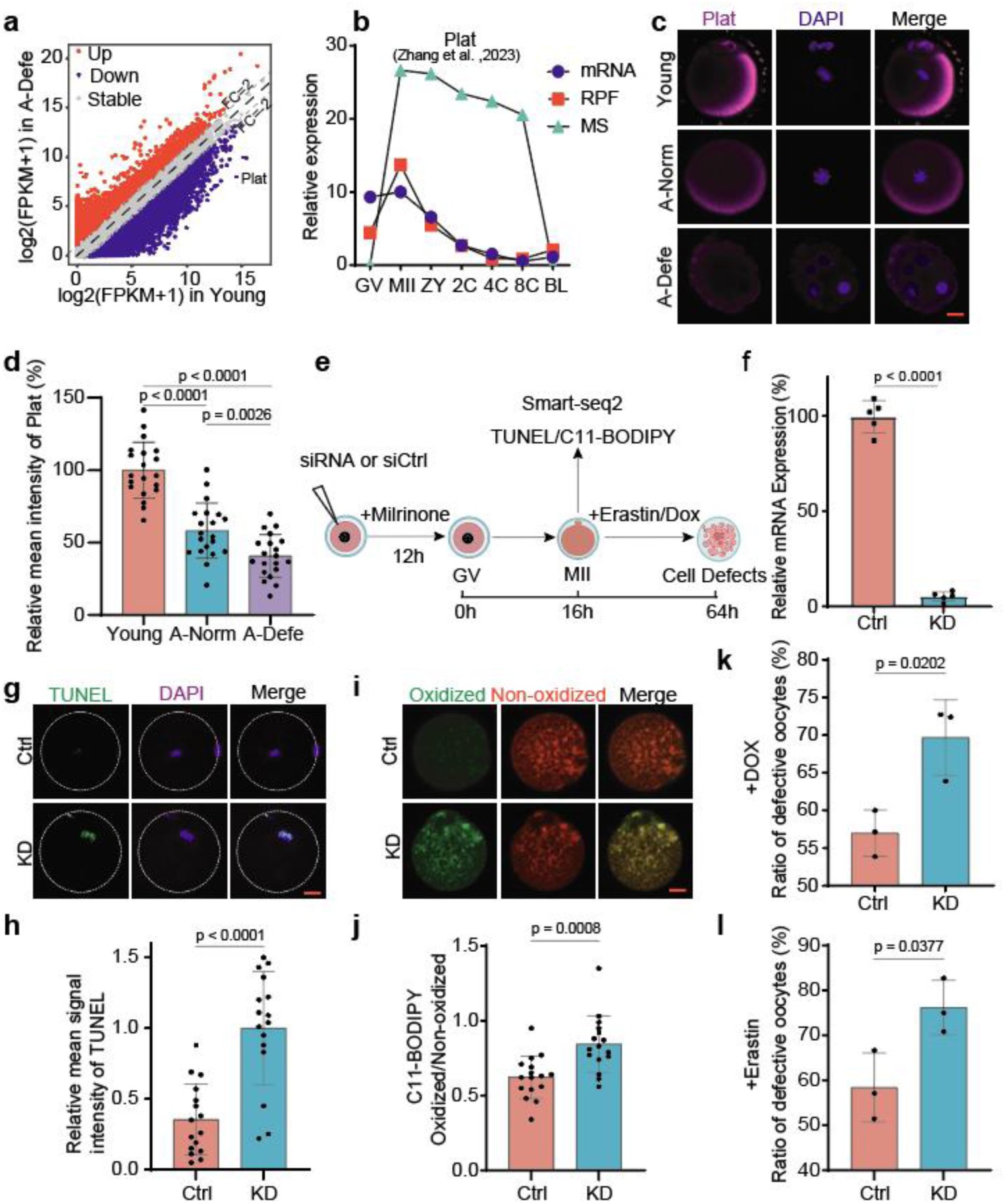
Reduced *Plat* expression in aged oocytes. (**A**) Violin plots comparing the expression of *Plat* in Young, A-Norm, and A-Defe oocytes. **** P < 0.0001 according to the Wald test implemented in DESeq2. (**B**) Manhattan plots demonstrating differential expression between MII- and GV-stage oocytes in the transcriptome, translatome, and proteome. Dots represent genes, and colors indicate different significant levels. (**C**) Venn diagram showing the overlaps between downregulated genes in A-Norm compared to Young oocytes and top upregulated proteins in GV-to-MII transition. The plot inside the overlapped region demonstrated that *Plat*-encoded protein was the top upregulated protein during the GV-to-MII transition among the 21 overlapped genes. (**D**) Representative images showing expression levels and distribution of Plat in Young, A-Norm, and A-Defe oocytes. Scale bar, 20 μm. (**E**) Barplots comparing relative fluorescence intensity of Plat between Young (n = 20), A-Norm (n = 20), and A-Defe (n = 20) oocytes. The data are represented as the mean ± s.d. ** P < 0.01 and **** P < 0.0001 according to two-tailed unpaired t-test. (**F**) Schematic diagram showing the workflow of the *Plat* knockdown experiments by siRNA. (**G**) Barplots showing the *Plat* knockdown efficiency at the mRNA levels by RT-PCR. *Gapdh* was used as an internal control. Experiments were replicated 5 times. The data are represented as the mean ± s.d. **** P < 0.0001 according to two-tailed unpaired t-test. (**H**) Representative images showing TUNEL levels in *Plat* knockdown (KD) and Control (Ctrl) oocytes. Scale bar, 20 μm. (**I**) Barplots comparing relative TUNEL fluorescence intensity in KD (n = 16) and Ctrl (n = 16) oocytes. The data are represented as the mean ± s.d. **** P < 0.0001 according to two-tailed unpaired t-test. (**J**) Representative images showing lipid peroxidation levels in *Plat* KD and Ctrl oocytes using C11-BODIPY staining (green for oxidize(D)red for non-oxidized). Scale bar, 20 μm. (**K**) Barplots comparing ratios of C11-BODIPY green to red fluorescence intensities in KD (n = 16) and Ctrl (n = 16) oocytes. The data are represented as the mean ± s.d. *** P < 0.001 according to two-tailed unpaired t-test. (**L**) Barplots comparing defect rates of young oocytes post-doxorubicin (DOX) treatment for 48h between KD (n = 98) and Ctrl (n = 97) oocytes. The data are represented as the mean ± s.d. * P < 0.05 according to two-tailed unpaired t-test. (**M**) Barplots comparing defect rates post Erastin treatment for 48h between KD (n = 112) and Ctrl (n = 93) oocytes. The data are represented as the mean ± s.d. * P < 0.05 according to two-tailed unpaired t-test.

While tPA is well known for its role in thrombus dissolution^39–41^, its function in mammalian oocytes remains poorly understood. To investigate this, we designed *Plat*-specific siRNA and introduced it into mouse GV-stage oocytes, followed by a 12-hour milrinone treatment to enhance siRNA efficacy (Fig. 5F). Upon oocyte maturation to the MII stage, qPCR confirmed efficient *Plat* silencing (Fig. 5G). In *Plat*-deficient MII oocytes, TUNEL assays revealed a significant increase in apoptosis (Fig. 5H,I), and lipid peroxidation assays indicated elevated ferroptosis levels (Fig. 5J,K), suggesting that *Plat* expression plays a crucial role in protecting oocytes from apoptotic and ferroptotic cell death. Furthermore, when treated with the apoptosis inducer Dox or the ferroptosis agonist Erastin, the *Plat*-deficient oocytes showed heightened susceptibility to both PCD pathways (Fig. 5L,M; Supplementary Fig. S4F,G).

In summary, combining results from transcriptomics and proteomics data analyses and *Plat*-deficiency experiments, we identified *Plat* as an essential maternal factor in MII oocytes, critical for preventing apoptosis and ferroptosis. Loss of *Plat* expression in aged oocytes appeared to drive PCD, contributing to their morphological defects.

### *Plat* activates the Erk1/2 pathway to prevent programed cell death in oocytes

To further investigate the mechanisms by which *Plat* prevents PCD in oocytes, we performed RNA-seq analysis on *Plat*-deficient oocytes and their controls. We identified 688 upregulated and 551 downregulated genes in *Plat*-deficient oocytes (Fig. 6A). Functional enrichment analysis revealed that the upregulated genes were primarily associated with apoptosis, while the downregulated genes were mostly linked to MAPK cascade and Ras signaling pathway (Fig. 6B). This aligns with earlier findings in morphologically defective oocytes, reaffirming the critical role of *Plat* in preventing oocyte PCD.

**Fig. 6.**
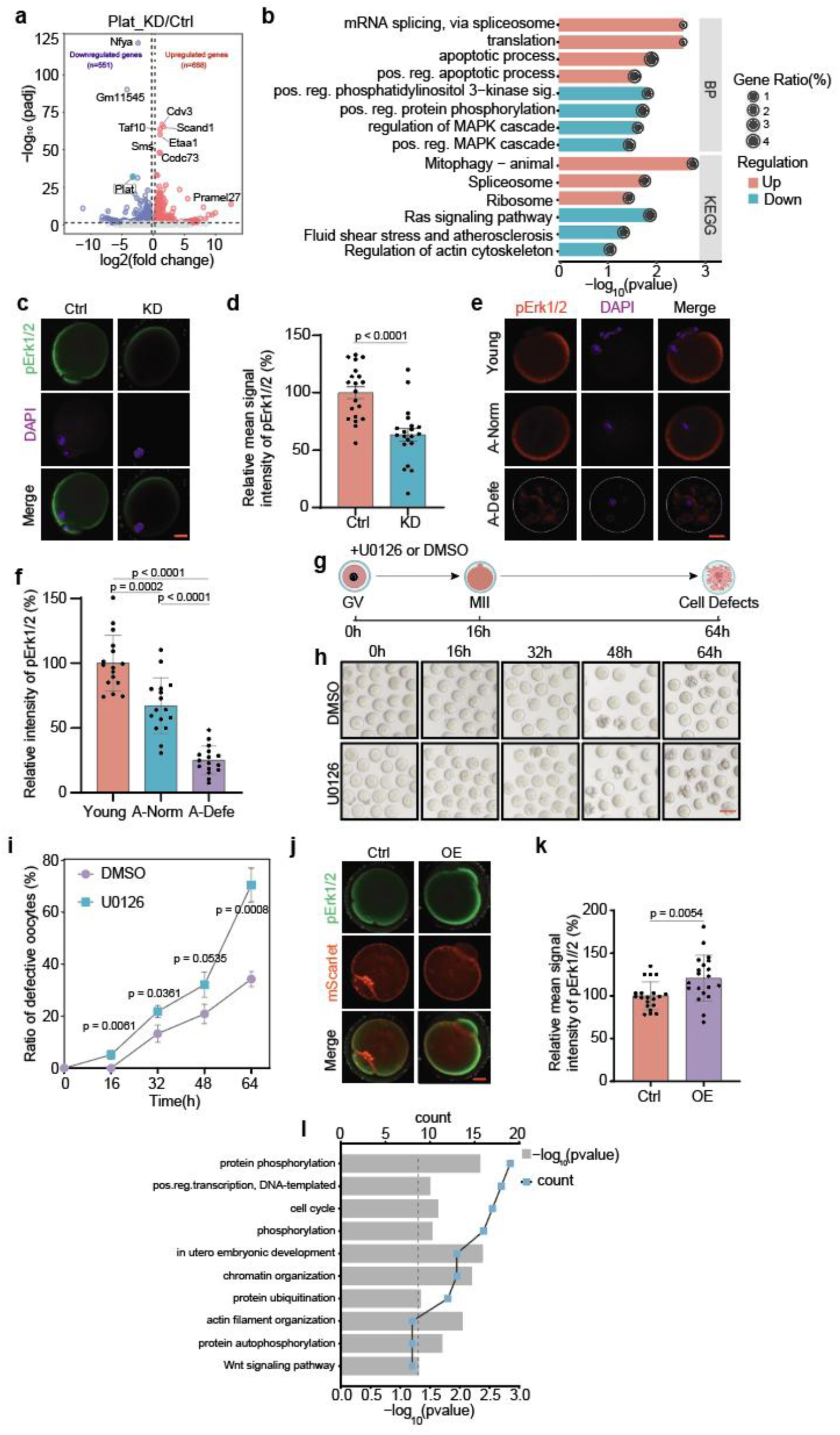
*Plat* affects oocyte survival through activating the Erk1/2 pathway. (**A**) Volcano plot showing differentially expressed genes (DEGs) following siRNA knockdown of *Plat*. Up and down-regulated DEGs were depicted in red and blue, respectively. The numbers of up- and down-regulated genes are indicated in the plot. (**B**) GO terms (biological processes) and KEGG pathways enriched in up- (red) or down-regulated (blue) genes in *Plat* knockdown oocytes. (**C**) Representative images showing pErk1/2 levels in *Plat* KD and Ctrl oocytes. Scale bar, 20 μm. (**D**) Barplots comparing relative fluorescence intensity of pErk1/2 in *Plat* KD (n = 20) and Ctrl (n = 20) oocytes. The data are represented as the mean ± s.d. **** P < 0.0001 according to two-tailed unpaired t-test. (**E**) Representative images showing pErk1/2 levels in Young, A-Norm, and A-Defe oocytes. DAPI-stained nuclei are shown in blue. Scale bar, 20 μm. (**F**) Barplots comparing relative fluorescence intensity of pErk1/2 in oocytes among Young (n = 16), A-Norm (n = 16), and A-Defe (n = 16) oocytes. The data are represented as the mean ± s.d. *** P < 0.001 and **** P < 0.0001 according to two-tailed unpaired t-test. (**G**) The scheme of U0126 treatment experiments for Erk1/2 activity inhibition. (**H**) Representative morphological images of oocytes treated with DMSO and U0126 at indicated time points. Scale bar, 100 μm. (**I**) Line plots comparing defect rates of oocytes treated with DMSO (n = 105) and U0126 (n = 96) at indicated time points. The data are represented as the mean ± s.d. * P < 0.05, ** P< 0.01 and *** P < 0.001 according to two-tailed unpaired t-test; ns, no significance. (**J**) Representative images showing the expression of mScarlet (red) and pErk1/2 (green) in *Plat* overexpression (OE) and Ctrl oocytes. Scale bar, 20 μm. (**K**) Barplots comparing relative fluorescence intensity of pErk1/2 between Ctrl (n = 20), and *Plat*-OE (n = 20) oocytes. The data are represented as the mean ± s.d. ** P < 0.01 according to two-tailed unpaired t-test. (**L**) Bar and line plots showing GO biological process terms enriched in 300 top-scored tPA-interacting proteins.

In *Plat*-deficient oocytes, the downregulated MAPK cascade and Ras signaling pathway converge on the Erk1/2 kinases^42^, which are pivotal regulators of cell survival^43^. Indeed, we observed inhibited Erk1/2 activation in *Plat*-deficient oocytes (Fig. 6C,D), analogue to the phenomenon observed in defective aged oocytes (Fig. 6E,F). To further validate the functional consequence of Erk1/2 inactivation in mouse oocytes, we treated GV-stage oocytes with U0126, a specific Erk1/2 inhibitor (Fig. 6G; Supplementary Fig. S5A). Our results demonstrated that Erk1/2 inhibition indeed led to significantly higher defect rates in oocytes, reaching more than twice that of controls after 64 hours (Fig. 6H,I). It has been reported that Erk1/2-knockout oocytes fail to develop beyond early embryonic stages due to fragmentation^44,45^. Reanalysis of the transcriptomic data from Erk1/2-knockout oocytes by Sha et al. revealed 3672 upregulated and 1288 downregulated genes (Supplementary Fig. S5B), associated with processes such as DNA damage response, apoptotic process, phosphorylation, and lipid metabolism. (Supplementary Fig. S5C,D). Detailed examination showed increased expression of apoptotic markers and ferroptosis drivers, along with decreased expression of ferroptosis suppressors in Erk1/2-knockout oocytes (Supplementary Fig. S5E–G), mirroring the apoptotic and ferroptotic changes observed in defective aged oocytes.

To determine the regulatory relationship between *Plat* expression and Erk1/2 activation, we microinjected in vitro transcribed *Plat* mRNA (Supplementary Fig. S5H) into GV-stage oocytes, in addition to performing the *Plat* deficiency experiments above (Fig. 6C,D). Overexpression of *Plat* significantly increased Erk1/2 activity in MII-stage oocytes (Fig. 6J,K). Interestingly, inhibition of Erk1/2 activity using U0126 did not decrease tPA levels but slightly increased them (Supplementary Fig. S5I), likely attributed to the negative feedback mechanisms within the Erk1/2 pathway^46^. The above findings suggest that *Plat* functions upstream of Erk1/2 activation.

To further explore how tPA regulates Erk1/2 activation, we conducted immunoprecipitation followed by mass spectrometry (IP-MS) to identify tPA-interacting proteins (Supplementary Fig. S6A). The 300 top-scored proteins were functionally involved in cytoskeletal organization and protein phosphorylation (Fig. 6L; Supplementary Fig. S6B). Notably, cytoskeletal proteins such as F-actin, Zp1-3, and keratin family members were enriched (Supplementary Fig. S6B), suggesting that tPA may contribute to maintaining oocyte structure. Indeed, F-actin intensity was significantly reduced in *Plat*-deficient oocytes, corroborating its role in cytoskeletal modulation (Supplementary Fig. S6C,D). More importantly, tPA was found to interact extensively with kinases involved in protein phosphorylation (Supplementary Fig. S6B). Among them, tyrosine kinases such as Alk and Txk are well-documented to induce Erk1/2 phosphorylation and activation^47,48^, suggesting that tPA regulates Erk1/2 activation through modulating key upstream regulators of the Erk1/2 pathway. In summary, our findings likely establish that *Plat* safeguards oocytes from PCD by activating the Erk1/2 pathway through phosphorylation, thus maintaining oocyte integrity and function.

### tPA supplementation ameliorates the quality of aged oocytes

Given that downregulation of *Plat* at the MII stage is one of the primary factors contributing to oocyte PCD, we next investigated whether supplementing tPA protein could counteract the reduced quality of aged oocytes. To this end, aged mouse GV oocytes were incubated in a medium enriched with exogenous tPA for 16 hours, and compared to young oocytes and aged oocytes without supplementation. As expected, tPA supplementation significantly improved the maturation rate of aged oocytes, reversing the observed decline in comparison to young oocytes (Fig. 7A,B). Moreover, tPA treatment markedly reduced the defect rate in aged oocytes, nearly restoring it to the level observed in young oocytes (Fig. 7C,D).

**Fig. 7.**
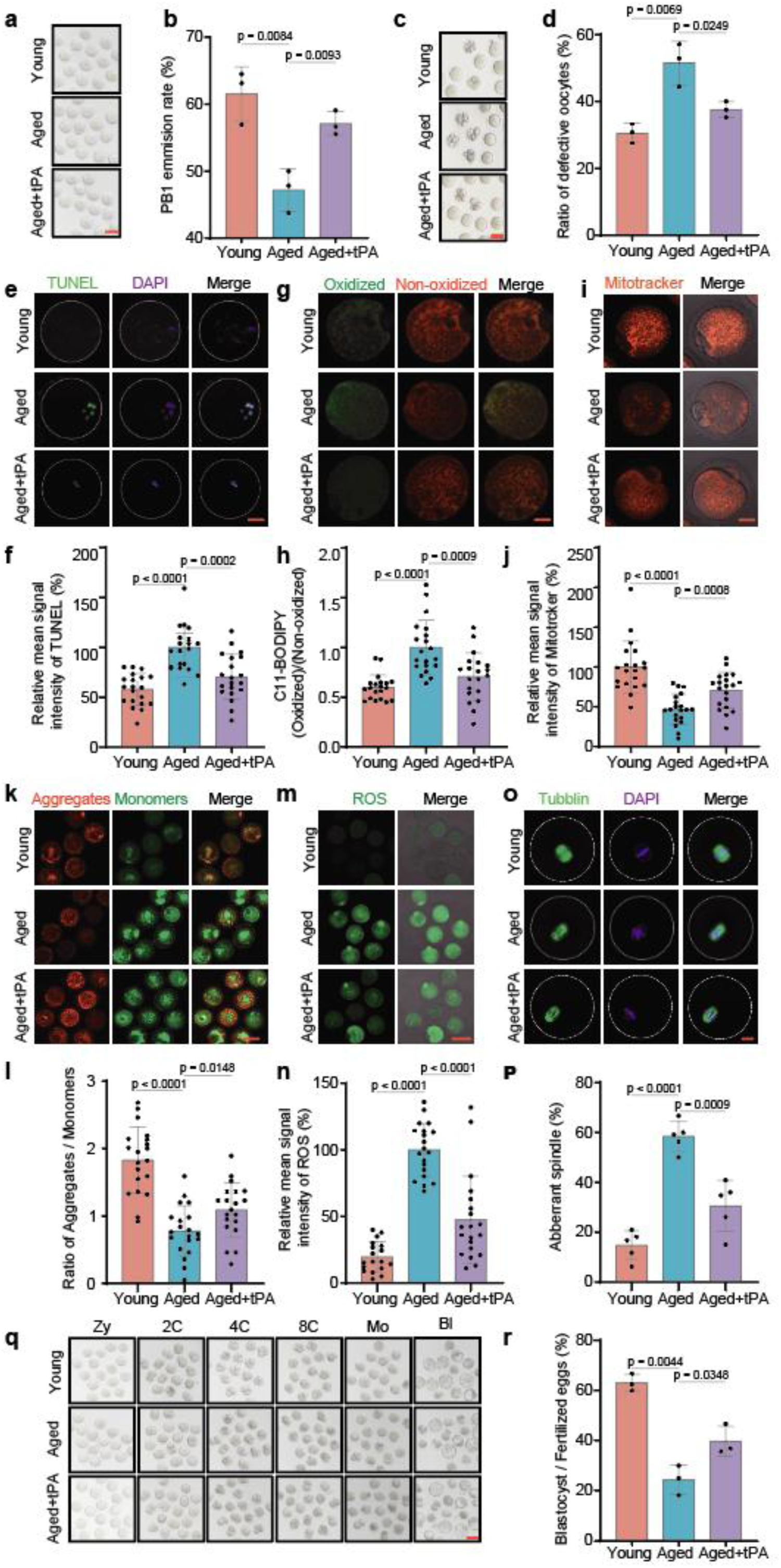
Exogenous tPA ameliorates the quality of aged mouse oocytes. (**A**) Representative images of in vitro matured oocytes in Young, aged, and tPA-treated aged (tPA+aged) groups. Scale bar, 100 μm. (**B**) Barplots comparing percentages of first polar body (PB1) extrusion between young (n = 191), aged (n = 85), and tPA+aged (n = 79) oocytes. The data are represented as the mean ± s.d. ** P< 0.01 according to two-tailed unpaired t-test. (**C**) Representative images of in vitro matured oocytes after culture for 48 hours in Young, aged, and tPA+aged groups. Scale bar, 100 μm. (**D**) Barplots comparing defect rates between young (n = 98), aged (n = 105), and tPA+aged (n = 106) oocytes. The data are represented as the mean ± s.d. * P < 0.05 and ** P< 0.01 according to two-tailed unpaired t-test. (**E**) Representative images showing TUNEL levels in oocytes of Young, aged, and tPA+aged groups. Scale bar, 20 μm. (**F**) Barplots comparing relative fluorescence intensity of TUNEL between young (n = 20), aged (n = 20), and tPA+aged (n = 20) oocytes. The data are represented as the mean ± s.d. *** P< 0.001 and **** P < 0.0001 according to two-tailed unpaired t-test. (**G**) Representative images showing lipid peroxidation levels in oocytes of Young, aged, and tPA+aged groups. The lipid peroxidation levels were assessed using C11-BODIPY (green, oxidized; red, non-oxidized). Scale bar, 20 μm. (**H**) Barplots comparing ratios of green to red fluorescence intensity in C11-BODIPY staining between young (n = 20), aged (n = 20), and tPA+aged (n = 20) oocytes. The data are represented as the mean ± s.d. *** P<0.001 and **** P < 0.0001 according to two-tailed unpaired t-test. (**I**) Representative images showing mitochondrial distribution in oocytes of Young, aged, and tPA+aged groups. Scale bar, 20 μm. (**J**) Barplots comparing relative mitochondrial fluorescence intensity in young (n = 19), aged (n = 20), and tPA+aged (n = 20) oocytes. The data are represented as the mean ± s.d. *** P< 0.001 and **** P < 0.0001 according to two-tailed unpaired t-test. (**K**) Representative images showing mitochondrial membrane potential (ΔΨm) in oocytes of Young, aged, and tPA+aged groups using JC-1 staining (red, high ΔΨm; green, low ΔΨm). Scale bar, 100 μm. (**L**) Barplots comparing ratios of red to green fluorescence intensity in JC-1 staining between young (n = 20), aged (n = 20), and tPA+aged (n = 20) oocyte. The data are represented as the mean ± s.d. * P< 0.05 and **** P < 0.0001 according to two-tailed unpaired t-test. (**M**) Representative images showing reactive oxygen species (ROS) levels in oocytes of Young, aged, and tPA+aged groups, measured by DCFH-DA staining. Scale bar, 20 μm. (**N**) Barplots comparing relative ROS fluorescence intensity between young (n = 20), aged (n = 20), and tPA+aged (n = 20) oocytes. The data are represented as the mean ± s.d. **** P < 0.0001 according to two-tailed unpaired t-test. (**O**) Representative images showing spindle morphology and chromosome alignment in oocytes of Young, aged, and tPA+aged groups after in vitro maturation. Scale bar, 20 μm. (**P**) Barplots comparing percentages of aberrant spindles between young (n = 131), aged (n = 153), and tPA+aged (n = 190) oocytes after in vitro maturation. The data are represented as the mean ± s.d. *** P< 0.001 and **** P < 0.0001 according to two-tailed unpaired t-test. (**Q**) Representative images of early embryos at indicated stages after Intracytoplasmic Sperm Injection (ICSI) on oocytes in Young, aged, and tPA+aged groups. Scale bar, 100 μm. ZY, zygote; 2C, 2-cell; 4C, 4-cell; 8C, 8-cell; Mo, morula; BL, blastocyst. (**R**), Barplots comparing blastocyst formation rate after ICSI on young (n = 43), aged (n = 37), and tPA+aged (n = 48) oocytes. The data are represented as the mean ± s.d. * P< 0.05 and ** P < 0.01 according to two-tailed unpaired t-test.

To determine whether the effects of tPA supplementation were mediated through mitigation of PCD, we performed TUNEL assays and lipid peroxidation staining. Both assays revealed significantly reduced apoptosis and lipid peroxidation levels in tPA-treated aged oocytes compared to untreated controls (Fig. 7E-H). These results directly demonstrated that tPA supplementation effectively protected aged oocytes from PCD. To further explore the mechanisms underlying tPA’s protective effects, we assessed mitochondrial function. MitoTracker staining indicated that tPA supplementation restored compromised mitochondrial activity in aged oocytes, and JC-1 staining demonstrated that mitochondrial membrane potential was significantly improved (Fig. 7I-L). Given the pivotal role of oxidative stress in apoptosis, we also examined reactive oxygen species (ROS) levels. As expected, tPA-treated aged oocytes exhibited significantly lower ROS levels than untreated aged oocytes (Fig. 7M,N). Given that tPA supplementation improved oocyte maturation (Fig. 7B), we further examined spindle morphology, which is a critical factor influencing oocyte maturation. Tubulin staining revealed that aged oocytes exhibited irregular spindle structures and chromosome misalignment, but tPA supplementation largely restored normal spindle morphology (Fig. 7O,P). These improvements in spindle integrity likely contributed to the enhanced maturation rate of tPA-supplemented aged oocytes.

Finally, to evaluate the developmental competence of tPA-treated aged oocytes, intracytoplasmic sperm injection (ICSI) was performed, followed by monitoring of embryonic development. Our data clearly demonstrated that the blastocyst formation rate, which was significantly reduced in untreated aged oocytes, was effectively restored with tPA supplementation (Fig. 7Q,R). Altogether, exogenous tPA supplementation markedly improves the quality of aged oocytes by restoring spindle morphology, enhancing mitochondrial function, reducing ROS levels, and subsequently mitigating apoptosis and ferroptosis. These effects collectively result in higher oocyte maturation rates, a reduced oocyte defect rate, and improved developmental potential, underscoring the therapeutic potential of tPA for age-related declines in oocyte quality.

## DISCUSSION

In addressing the mechanisms underlying the decline in oocyte quality during female reproductive aging, we integrate soRNA-seq analysis and functional research in this study, revealing that morphologically defective oocytes from aged mice, devoid of developmental potential, are subject to apoptotic and ferroptotic death. These processes are generally suppressed in MII-stage oocytes of young mice with high *Plat* expression. This is because tPA, encoded by *Plat*, interacts with phosphorylation kinases to activates the Erk1/2 pathway, which has been shown to be essential for oocyte maturation and further development into embryos^44,49^. Conversely, the downregulation of *Plat* in defective oocytes emerges as a critical molecular event exacerbating PCD and contributing to oocyte degeneration. These findings provide novel insights into the molecular underpinnings of oocyte aging, highlighting *Plat* as a key regulator of oocyte viability. Moreover, our successful rescue of defective oocytes through exogenous tPA supplementation opens new avenues to enhance reproductive outcomes for women of advanced maternal age.

In this study, we included oocytes with abnormal morphologies for the initial analysis, which is different from typical studies on oocyte aging, where defective oocytes were usually discarded beforehand because they were not considered clinically rescuable. Aiming at finding out what events have occurred following oocyte degeneration, we considered the defective oocytes as a final stage of oocyte degradation. Thanks to this experiment design, we have actually found out that aged oocytes are more susceptible to both apoptosis and ferroptosis, and *Plat* loss in aged oocytes is likely the key to trigger oocyte-specific PCD. Furthermore, we utilized the advanced soRNA-seq approach to profile transcriptomes of individual oocytes, which enables us discarding defective oocytes in downstream analysis, such as comparisons between young oocytes and aged oocytes with normal morphology. These data would help us to dissect the degeneration process of aged oocytes at different stages, hence facilitating mechanistic discoveries.

Our study highlights the complexity of PCD in oocytes, particularly the co-occurrence of apoptosis and ferroptosis in defective oocytes. While previous studies have predominantly focused on apoptotic pathways^50^, our findings expand this understanding by implicating ferroptosis through evidence of increased lipid peroxidation and disrupted iron metabolism in defective oocytes. Notably, while playing a pivotal role in granulosa cell apoptosis^51^, the absence of Caspase-3 expression in full-grown oocytes underscores its limited role in oocyte apoptosis. Therefore, PCD in aged oocytes may result from a synergistic involvement of both pathways, a hypothesis supported by the apoptosis and ferroptosis dual inhibition experiments (Fig. 4K). Nonetheless, other forms of PCD, such as necroptosis^52^ or autophagy-dependent cell death^53^, may also contribute to oocyte degeneration, necessitating further investigation into their roles and interactions.

tPA, known as tissue plasminogen activator, was initially identified as a fibrinolytic enzyme^54,55^. Its non-fibrinolytic roles in neuronal protection and cellular survival have been increasingly recognized^56,57^. For example, Lee et al have revealed the pivotal role of tPA in promoting the proliferation of neuronal cells^58^, and studies by Hua et al. and Lin et al. demonstrate its anti-apoptotic effects in granulosa cells and oviductal epithelial cells^59^, and macrophages^60^, respectively. Our study extends its function to oocytes, identifying tPA as a maternal factor crucial for supressing both apoptosis and ferroptosis at the MII stage, underlining its regulatory roles in dual cell death pathways. In addition, tPA may play distinct roles at different stages of oocyte development or maturation, because one previous study has demonstrated that its overexpression in GV-stage bovine cumulus–oocyte complexes impairs oocyte maturation^61^. The shift of tPA from a harmful to a beneficial role also explains its sharp protein-level increase during the transition from the GV stage to the MII stage^32,38^, which points to intricate post-transcriptional regulation, as its RNA expression at both stages does not change obviously (Fig. 5B). This observation highlights that post-transcriptional regulation, particularly translational activation, is critically important during oocyte meiotic maturation, and such timely protein availability might be attributed to mRNA’s poly(A) tail elongation or 3’UTR isoform switches during this transition, as we previously reported^62^.

Erk1/2 inactivation in *Plat*-deficient oocytes mirrors the defects observed in aged oocytes, but tPA expression remains largely unchanged in Erk1/2-inhibited oocytes, suggesting that *Plat* operates upstream of Erk1/2. In addition, our IP-MS experiment reveals that tPA interacts with tyrosine kinases implicated in phosphorylation, which may subsequently activate the Erk1/2 pathway, reinforcing a regulatory connection between tPA expression and Erk1/2 activation. Activated Erk1/2 enhances mitochondrial stability and suppresses pro-apoptotic signals, thereby curbing the intrinsic apoptotic pathway^63,64^. Similarly, Erk1/2-mediated transcriptional regulation of antioxidant genes contribute to maintaining redox homeostasis, thereby limiting lipid peroxidation, a key driver of ferroptosis^65,66^. In oocytes, Erk1/2 inactivation correlates with elevated apoptosis and ferroptosis markers, underscoring its dual protective role against the both PCD forms. Future studies should explore the mechanistic intricacies of Plat-Erk1/2 regulation and its interaction with other PCD pathways to fully elucidate its role in maintaining oocyte health. Furthermore, the identification of cytoskeletal proteins among tPA-interacting partners highlights additional roles in preserving oocyte structure and function.

In conclusion, we provide a mechanistic basis for the decline in oocyte quality with aging, attributed to reduced *Plat* expression and downstream Erk1/2 inactivation. The successful restoration of oocyte quality via exogenous tPA supplementation highlights its potential clinical application. This strategy aligns with the growing optimism to improve the fertility prospects of reproductive aged women through systemic interventions, such as dietary supplementation. As the U.S. Food and Drug Administration (FDA)-approved agent for ischemic stroke^67^, tPA’s safety profile in humans is well-established and thus underscores its therapeutic promise, as long as coagulation-related risks are managed.

## MATERIALS AND METHODS

### Mice

All female ICR mice were obtained from the Experimental Animal Center of Nanjing Medical University. They were housed under controlled environmental conditions in a Specific Pathogen Free (SPF) setting, with a temperature of 20–22 °C, a light/dark cycle of 12/12 hours, and humidity of 50– 70%. Food and water were provided ad libitum. All experimental protocols involving mice were approved by the Ethics Committee of Nanjing Medical University (No.: 2410009), with care and use following applicable guidelines and regulations.

### Oocyte collection

For the collection of germinal vesicle (GV) stage oocytes, female mice aged 3-8 weeks (young) or 10-12 months (aged) were intraperitoneally injected with 5 IU or 10 IU of pregnant mare’s serum gonadotropin (PMSG) (Ningbo Sansheng Pharmaceutical, China). After 48 h, the mice were humanely sacrificed. The ovaries were harvested and the extraneous fat tissue was removed. The oocytes were released by mincing the ovaries, using a blade, in manipulation medium and collected with a handmade glass capillary of approximately 150 micrometers in diameter. The collected oocytes were cultured in M2 medium (Nanjing Luanchuang Co., China) for 16 h at 37 ℃ and 5% CO_2_, with preference given to MII oocytes that had extruded their first polar body for subsequent experiments. For the collection of MII oocytes, young and aged mice were injected separately with 5 IU or 10 IU PMSG. 46 h after injection, they received injections of 5 IU or 10 IU human chorionic gonadotropin (HCG) (Ningbo Sansheng Pharmaceutical, China). After 14-16 h, the mice were sacrificed and the cumulus-oocyte complexes were collected from the ampulla of the fallopian tube. The complexes were incubated in M2 medium at 37 ℃ and treated with 0.3% hyaluronidase to remove cumulus cells and retrieve clean oocytes.

### in vitro fertilization (IVF) and embryo collection

Female mice were intraperitoneally injected with 5 IU (young) or 10 IU (aged) PMSG. At 46-48 h after PMSG injection, 10 IU or 5 IU HCG was injected to induce superovulation. Female mice were euthanized by cervical dislocation ∼16 h after HCG injection, and cumulous oocyte complexes (COCs) were isolated from oviducts in HTF medium drops submerged in mineral oil. Sperms were isolated from cauda epididymi and pre-incubated in TYH (Nanjing Luanchuang Co., China). Following recommended incubation, sperm were added to COC drops and incubated (37°C, 5% CO2 in air) for ∼3–5 h to allow for fertilization to occur and subjected to a further incubation of ∼4–6 h. At this time, cumulus cells detached from the zygotes, which were further washed in drops of KSOM media (Nanjing Luanchuang Co., China), with any degenerating zygotes being discarded. Viable zygotes were transferred to drops of KSOM media overlaid with mineral oil for embryo collection. Embryos were harvested at specific time intervals after injection of HCG: 27-29 hours (zygotes), 33–35 hours (early 2-cell), 46–48 hours (late 2-cell), 62–65 hours (8-cell) and 94–96 hours (blastocyst).

### Western blot

For immunoblotting, 100 oocytes were quickly rinsed three times with phosphate-buffered saline (PBS) and then lysed for 10 min in RIPA (YEASEN, 20101ES60) containing 1% protease inhibitors (CWBIO, CW2200S) and 1% phosphatase inhibitors (CWBIO, CW2383S). Samples were heated at 95 °C for 5 min and separated on precast polyacrylamide gels (GenScript, M00929) using running buffer. Proteins were then transferred onto a polyvinylidene fluoride (PVDF) membrane (GVS, 1214429) and blocked with blocking buffer (ABclonal, RM02956) for 30 min at room temperature before being incubated with primary antibodies overnight at 4 ℃. After washing, membranes were incubated with HRP-conjugated secondary antibodies for 1 h at room temperature. An enhanced chemiluminescence (ECL) substrate (UE, S6010L) was used for signal detection via an automated chemiluminescence imaging system.

### Immunofluorescence

Mice oocytes were rinsed with PBS and fixed with 4% paraformaldehyde for 30 min, followed by washing with a buffer containing Tween 20 and Triton X-100. Permeabilization with Triton X-100 in PBS was performed for 20 min at room temperature. Samples were blocked with 1% BSA in PBS for 1 h and incubated with primary antibodies overnight at 4 ℃. After washing, samples were incubated with the appropriate secondary antibodies conjugated with fluorophores at room temperature for 1 h. DAPI was used for counterstaining. Samples were then mounted on glass slides and examined using a confocal laser scanning microscope. Fluorescence intensities were measured using the ImageJ software.

### Antibodies and reagents

Antibodies used in this study include Anti-Plat antibody (zenbio, R381605), anti-cleaved Caspase8 antibody (zenbio, 250106), anti-Bax antibody (ABclonal, A12009), anti-Caspase3 antibody (ABclonal, A19654), anti-Gpx4 antibody (zenbio, 381958), anti-pErk1/2 antibody (ABclonal, A18196), anti-Gapdh antibody (proteintech, 60004-1-Ig), anti-Ddb1 antibody (ABclonal, A2896), and anti-β-tubulin antibody (ABclonal, A12289). The aforementioned antibodies were diluted at 1:200 for immunofluorescence and at 1:500 for western blot. Horseradish peroxidase (HRP)-conjugated goat anti-rabbit IgG (H+L) (ABclonal, AS014), diluted at 1:3000, HRP-linked goat anti-mouse IgG (H+L) antibody (ABclonal, AS003), diluted at 1:3000, CoraLite488-conjugated goat anti-rabbit IgG (H+L) (proteintech, SA00013-2), diluted at 1:250, CoraLite594-conjugated goat anti-rabbit IgG (H+L) (proteintech, SA00013-4), diluted at 1:250, CoraLite488-conjugated goat anti-mouse IgG (H+L) (proteintech, SA00013-1), diluted at 1:250, CoraLite594-conjugated goat anti-mouse IgG (H+L) (proteintech, SA00013-3), diluted at 1:250.

### Construction of *Plat* recombinant vector

The *Plat* cDNA was amplified using Platinum SuperFi II PCR Master Mix (Invitrogen, 12368010) and ligated into a T7-driven plasmid vector using a ClonExpress II One Step Cloning Kit (Vazyme, C112-01). The recombinant plasmid was transformed into competent E. coli DH5a (Vazyme, C502-02) and the correct construction was confirmed by Sanger sequencing.

### in vitro transcription

For the preparation of mRNA for microinjection, the plasmid was digested to be linearized (NEB, R0560V), followed by gel purification. HiScribe® T7 ARCA mRNA Kit (with tailing) (NEB, E2060S) was used for in vitro transcription and tailing of the linearized DNA, which was then recovered by lithium chloride precipitation and resuspended in nuclease-free water.

### Oocyte microinjection

Milirinone was added to M2 culture medium to inhibit spontaneous germinal vesicle breakdown (GVBD). All microinjections were performed using a micromanipulation system. mRNA or siRNA was injected into the oocytes. The concentration of the injected mRNA was 500 ng/µL, and siRNA was at a concentration of 20μM. The siRNA targeting *Plat* and the negative control were designed and synthesized by Tsingke. After injection, oocytes were cultured in M16 medium with 2μM milirinone for 10-20 h to ensure the mRNA or siRNA took effect, then transferred to M16 without milirinone for continued culture.

### Determination of mitochondrial membrane potential

The mitochondrial membrane potential was assayed using a kit containing JC-1 (Beyotime, C2006). Briefly, oocytes were stained in M2 medium with JC-1 for 30 min at 37 °C, washed with JC-1 buffer, and examined under the confocal laser scanning microscope (Zeiss LSM 900, Carl Zeiss AG, Germany).

### Detection of divalent iron ions

Oocytes were stained for 30 min in HBSS solution containing 1μM divalent iron ion detection probes (DOJINDO, F374) as per the manufacturer’s instructions and examined under the confocal laser scanning microscope (Zeiss LSM 900, Carl Zeiss AG, Germany).

### Detection of lipid peroxidation

Lipid peroxidation was detected using the Image-iT Lipid Peroxidation Kit (Thermofisher, C10445). Briefly, 50 oocytes were incubated in M2 medium containing 10μM Lipid Peroxidation Sensor for 30 min at 37 °C, washed three times with M2, and examined under the confocal laser scanning microscope (Zeiss LSM 900, Carl Zeiss AG, Germany).

### Mitochondria labeling with MitoTracker

For mitochondrial visualization, oocytes were incubated in a diluted solution of MitoTracker™ Deep Red dye (Thermofisher, M22426) and examined under a confocal laser scanning microscope.

### Detection of reactive oxygen species (ROS)

ROS were detected using an assay kit. Oocytes were stained in M16 culture medium containing 2,7-Dichlorodi -hydrofluorescein diacetate (DCFH-DA), washed, and examined under a confocal laser scanning microscope.

### RT-PCR

Twenty oocytes per sample were lysed in 2 μl of lysis buffer (0.2% Triton X-100: RNase inhibitor = 9:1) and reverse transcription was performed using the HiScript III 1st Strand cDNA Synthesis Kit (+gDNA wiper) (Vazyme, R312-01). Real-time RT-PCR analysis was conducted using PerfectStart Green qPCR SuperMix (Transgen, AQ602-11) and the QuantStudio 3 real-time PCR system (Thermofisher). Data were calculated using the 2^-ΔΔCt^ method with Gapdh as the reference.

### Immunoprecipitation followed by mass spectrometry (IP-MS)

We conducted IP experiments utilizing the Pierce Classic Magnetic IP/Co-IP Kit (ThermoFisher, 88804). Initially, 1500 MII oocytes were collected and lysed in IP lysis buffer. The cell lysate was incubated overnight at 4°C with antibody. Subsequently, the antigen-antibody complex was coupled with protein A/G magnetic beads for 1 h at room temperature, followed by two washes with IP lysis/wash buffer and a final wash with pure water. The samples were then heated for 10 min at 95°C after the addition of SDS, and the separated proteins were resolved by SDS-PAGE. LC-MS/MS analysis was performed by Luming Biology (Shanghai, China). Briefly, the examined protein bands were subjected to enzymatic digestion (0.02μg/μL trypsin in digestion buffer) and peptide desalting to form peptide samples for further analysis. The analytical equipment comprised an Ultimate 3000nano ultra-high-performance liquid chromatography tandem with Q Exactive plus high-resolution mass spectrometer (ThermoFisher Scientific). The UltiMate 3000 RSLCnano Nano-HPLC system (ThermoFisher Scientific) was used for sample separation. The mobile phase A liquid was a 0.1% formic acid in water solution, while B liquid contained 0.1% formic acid in acetonitrile. The trap column, 100 μm × 20 mm (RP-C18, Agilent) was balanced with 100% A liquid at a flow rate of 3 μL/min. Samples were loaded onto the trap column by an autosampler and then resolved through an analysis column, 75 μm × 150 mm (RP-C18, New Objective, USA) with a flow rate of 300 nL/min. Blank solvent was used to rinse between samples for 30 min of gradient washing. The enzymatic digests were separated by capillary high-performance liquid chromatography followed by mass spectrometry analysis using the Q-Exactive Plus mass spectrometer. Detection settings were calibrated with standard liquid before use, with parent ion scan range: 300-1500 m/z. The mass spectrometry scanning method was Data Dependent Acquisition (DDA), where 20 most intense fragment spectra (MS2 scan) were collected after each full scan, with fragmentation performed by high-energy collision dissociation (HCD), NCE energy set at 28, and dynamic exclusion time at 25 sec. The MS1 resolution was set at 70,000 with m/z 200, AGC target at 3e6, maximum injection time at 100 ms. For MS2, the resolution was set at 17,500, the AGC target at 1e5, and the maximum injection time at 50 ms. The sample was subject to protein identification (relative quantification) analysis using the database searching software ProteomeDiscover 2.5. The obtained protein list reflects the types and relative quantities of proteins detected in this mass spectrometry examination.

### Intra-cytoplasmic sperm injection (ICSI)

Young (3-8 weeks) and aged (10-12 months) ICR mice were intraperitoneally injected with 5 or 10 IU of PMSG. After 48 h, GV stage oocytes were collected and cultured in M2 medium with or without tPA. After 16 h, cells that had expelled PB1 were selected. A 10-week-old ICR male mouse was humanely euthanized, and the sperm from the epididymis was harvested and released into HTF medium. Subsequently, the sperm and oocytes were transferred to CZB culture medium, and a piezoelectrically driven micromanipulator was employed to inject the sperm head into the oocytes. All oocytes were then further cultured in a KSOM medium at 37 °C with 5% CO2 until reaching the blastocyst stage.

### RNA-Seq library preparation and sequencing

RNA-seq libraries were constructed following the Smart-seq2 protocol ^68^. In brief, the AccuNext Single Cell/Low Input cDNA Synthesis & Amplification kit (Accurate Biology, AG12501) was used for reverse transcription and full-length amplification. Individual oocytes or ten-oocyte samples were lysed in a 2 μl lysis buffer (0.2% Triton X-100 mixed with RNase inhibitor in a 9:1 ratio). After adding 3’ Oligo (dT) Primer, the samples were incubated at 72°C for 3 min, followed by cooling on ice. Prepared RT Master Mix was then processed on a PCR machine with the following cycles: 42°C for 90 min, 70°C for 10 min, with a hold at 4°C. The amplified products were mixed with 2X AccuNext PCR Buffer and PCR Primers, followed by 16 cycles of amplification. The TransNGS® Tn5 DNA Library Prep Kit for Illumina (for 1 ng DNA) (Transgen, KP111-03) was used for DNA fragmentation and the addition of sequencing adapters. Fragments were size-selected using 0.6X and 0.15X DNA Clean Beads (Vazyme, N411-01). Finally, the library was sequenced on the Illumina NovaSeq 6000.

### RNA-seq data analyses

For upstream data processing, Trim Galore (v0.6.6)^69^ or flexbar (v2.5) was employed to remove low-quality bases and adaptors. Subsequently, the reads were mapped to the reference mouse genome (mm10) using the STAR (v2.7.10b) aligner^70^. HTSeq (v0.13.5)^71^ was used to generated sample-by-gene read count matrix. The workflow for single-oocyte data processing is conducted according to the descriptions in previous publication ^72^. In brief, file operations were conducted in R (version 4.0.4), and Seurat package (v4.1.1)^73^ was mainly used for analysis, including read count normalization, highly variable gene identification, expression data scaling, and principal component analysis (PCA). The number of PCs for downstream analysis was determined using the JackStraw method, based on which UMAP was conducted. The R package DESeq2 (v1.30.1)^74^ was used for differential gene expression analysis. Genes with adjusted P value < 0.05 and log2-transformed fold-change > 1.2 or < –1.2 were considered as differentially expressed. DAVID online tool (version 2021)^75^ was used for GO enrichment analysis. If the number of differentially expressed genes > 500, then the top 500 were taken for GO analysis. For metabolic flux analysis, we utilized the commands provided by the COMPASS software package (v0.9.10.2)^35^.

### Statistical analysis

No statistical methods were used to predetermine sample sizes, but our sample sizes are similar to those reported in previous publications^76,77^. Statistics were calculated using GraphPad Prism (version 9.0.0) software or R software (version 4.0.4). All experiments were replicated more than three times, and the data obtained were subjected to statistical analysis. Data are presented as arithmetic mean ± s.d. unless otherwise indicated in the figure legends. Differences between two groups were analyzed for statistical significance using two-tailed unpaired Student’s t-tests or two-tailed Fisher’s exact test unless otherwise indicated. P < 0.05 was considered to be significant unless otherwise stated. For the boxplots in the figures, center bars indicate median of the data, the box edges represent the third and the first quartiles, and the whiskers show the 1.5× interquartile range. For the violin plot, the shaded area represents the data distribution. Sample sizes or numbers (*n* value) of independent experiments are indicated in the figure legends.

## ETHICS DECLARATION

All methods were performed in accordance with the relevant guidelines and regulations. Animal (mice) procedures were approved by the Ethics Committee of Nanjing Medical University (No.: 2410009), with care and use following applicable guidelines and regulations.

## AUTHOR CONTRIBUTIONS

Xi Wang, Yuanlin He and Xingsi He conceived of the project. Xi Wang, Xingsi He, Hanwen Zhang, Ya Wang, and Yuanlin He designed the experiments. Xingsi He and Hanwen Zhang performed the experiments with help from Huanyu Yan, Qiaozhen Shi and Xiao Zeng. Xingsi He and Hanwen Zhang analyzed the data with help from Qiuzhen Chen, Min Su, Wei Shen, Yangmin Wang, Chikun Wang, and Shuyue Hou. Xingsi He and Xi Wang drafted the manuscript. Xi Wang, Xingsi He, Yuanlin He and Zhibin Hu reviewed and edited the manuscript. All authors read and approved the manuscript.

## ACKNOWLEDGEMENTS

We would like to express our sincere gratitude to Prof. Chaojun Li and Prof. Ran Huo for their constructive suggestions on this work. This study was partially supported by the National Natural Science Foundation of China (32170742 and 92374117 to X.W.), the National Key Research & Development (R&D) Program of China (2022YFC2702500 to X.W.), and the Natural Science Foundation of Jiangsu Province (BK20220201 to Y.W.).

