## supplementary figures for "*Plat* safeguards maternally aged oocytes against programmed cell death through activating the Erk1/2 pathway" (from another version of this preprint: version 2, 2025-02-12)

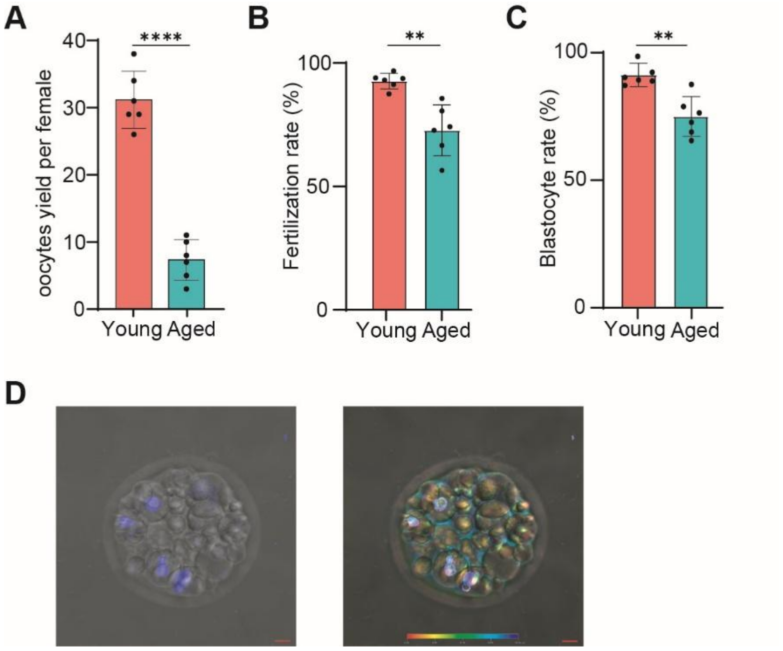


**Supplementary Fig. S1 | 3D morphology of defective oocytes.** (**A**) The number of oocytes extruded by the superovulation procedure in aged mice (n=6) was significantly lower than that of young mice (n=6). The data are presented as the mean ± s.d. from 6 biologically independent samples. The data are represented as the mean ± s.d. **** P < 0.0001 according to two-tailed unpaired t-test. (**B**) The fertilization rate of oocytes from aged mice (n=6) was significantly lower than that of young mice (n=6) following in vitro fertilization (IVF). The data are presented as the mean ± s.d. from 6 biologically independent samples. The data are represented as the mean ± s.d. ** P< 0.01 according to two-tailed unpaired t-test. (**C**) The blastocyst rate of oocytes from aged mice (n=6) was significantly lower than that of young mice (n=6). Data represented as the mean ± s.d. from 6 biological replicates. The data are represented as the mean ± s.d. ** P< 0.01 according to two-tailed unpaired t-test. (**D**) Left: Bright-field image depicting defective oocytes from aged mice. Right: Depth-encoded visualization of the same defective oocytes from aged mice. The DAPI stain is indicated in blue. Scale bar: 10μm.

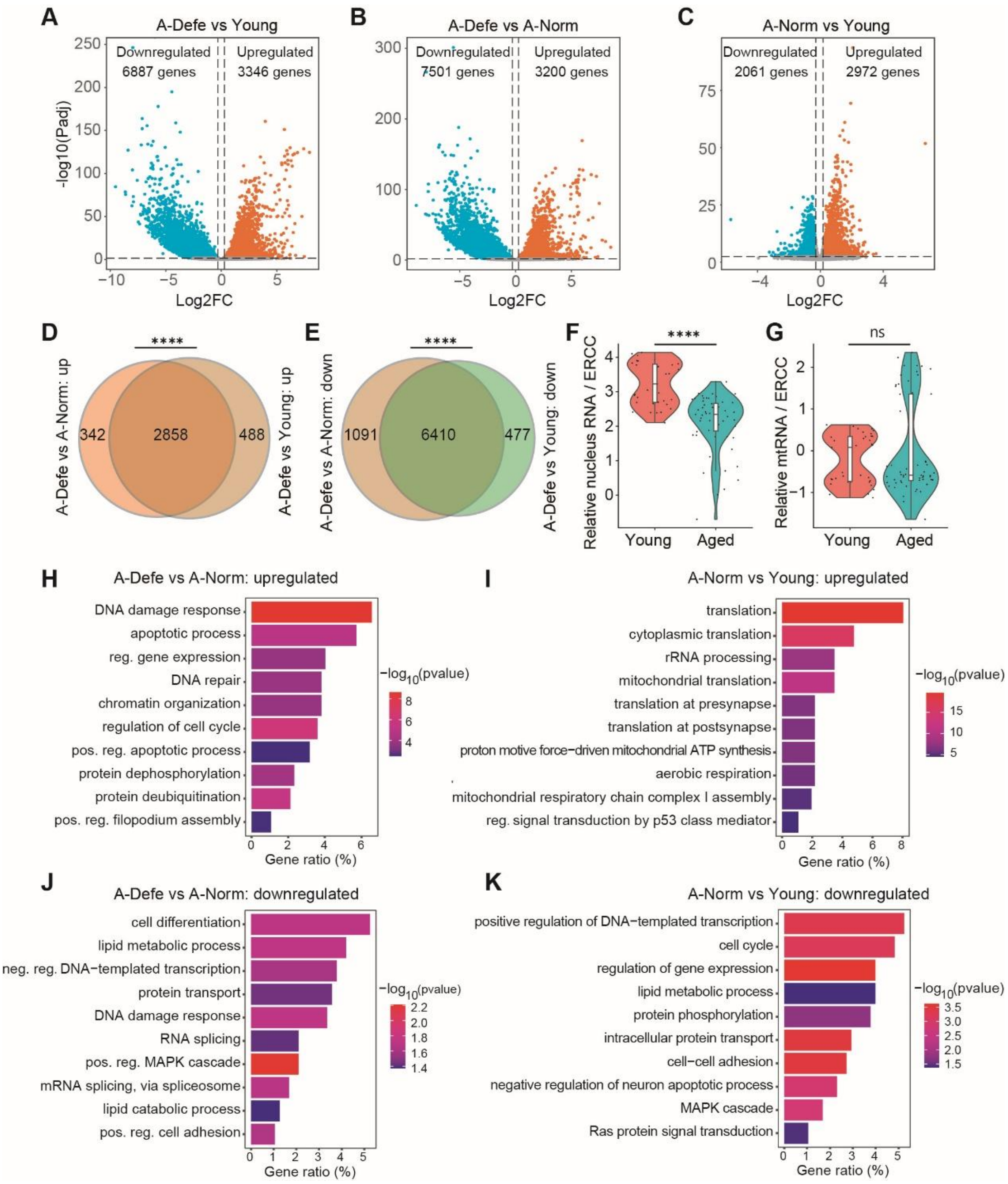


**Supplementary Fig. S2 | Differential gene analysis between oocyte clusters.** (**A**) Volcano plot showing up- (red) and down-regulated (blue) genes between Aged-Defe and Young oocytes. The numbers of up- and down-regulated genes are indicated in the plot. (**B**) Volcano plot showing up- (red) and down-regulated (blue) genes between Aged-Defe and Age-Norm oocytes. The numbers of up- and down-regulated genes are indicated in the plot. (**C**) Volcano plot showing up- (red) and down-regulated (blue) genes between Aged-Norm and Young oocytes. The numbers of up- and down-regulated genes are indicated in the plot. (**D**) Venn diagram illustrating the overlap of upregulated genes in comparisons between A-Defe and A-

Norm, and between A-Defe and Young, with specific gene numbers indicated in the diagram. **** $P < 0.0001$ according to Fisher's exact test. (**E**) Venn diagram illustrating the overlap of downregulated genes in comparisons between A-Defe and A-Norm, and between A-Defe and Young, with specific gene numbers indicated in the diagram. **** $P < 0.0001$ according to Fisher's exact test. (**F**) Violin plots comparing ratios of nucleus RNA expression relative to ERCC RNA between young and aged oocytes. **** $P< 0.0001$ according to two-tailed unpaired t-test. (**G**) Violin plots comparing ratios of mtRNA expression relative to ERCC RNA between young and aged oocytes. Statistical significance was determined by two-tailed unpaired t-test. ns, no significance. (**H**) Enriched GO terms in the top 500 upregulated genes between A-Defe and A-Norm oocytes. (**I**) Enriched GO terms in the top 500 upregulated genes between A-Norm and Young oocytes. (**J**) Enriched GO terms in the top 500 downregulated genes between A-Defe and A-Norm oocytes. (**K**) Enriched GO terms in the top 500 downregulated genes between A-Norm and Young oocytes.

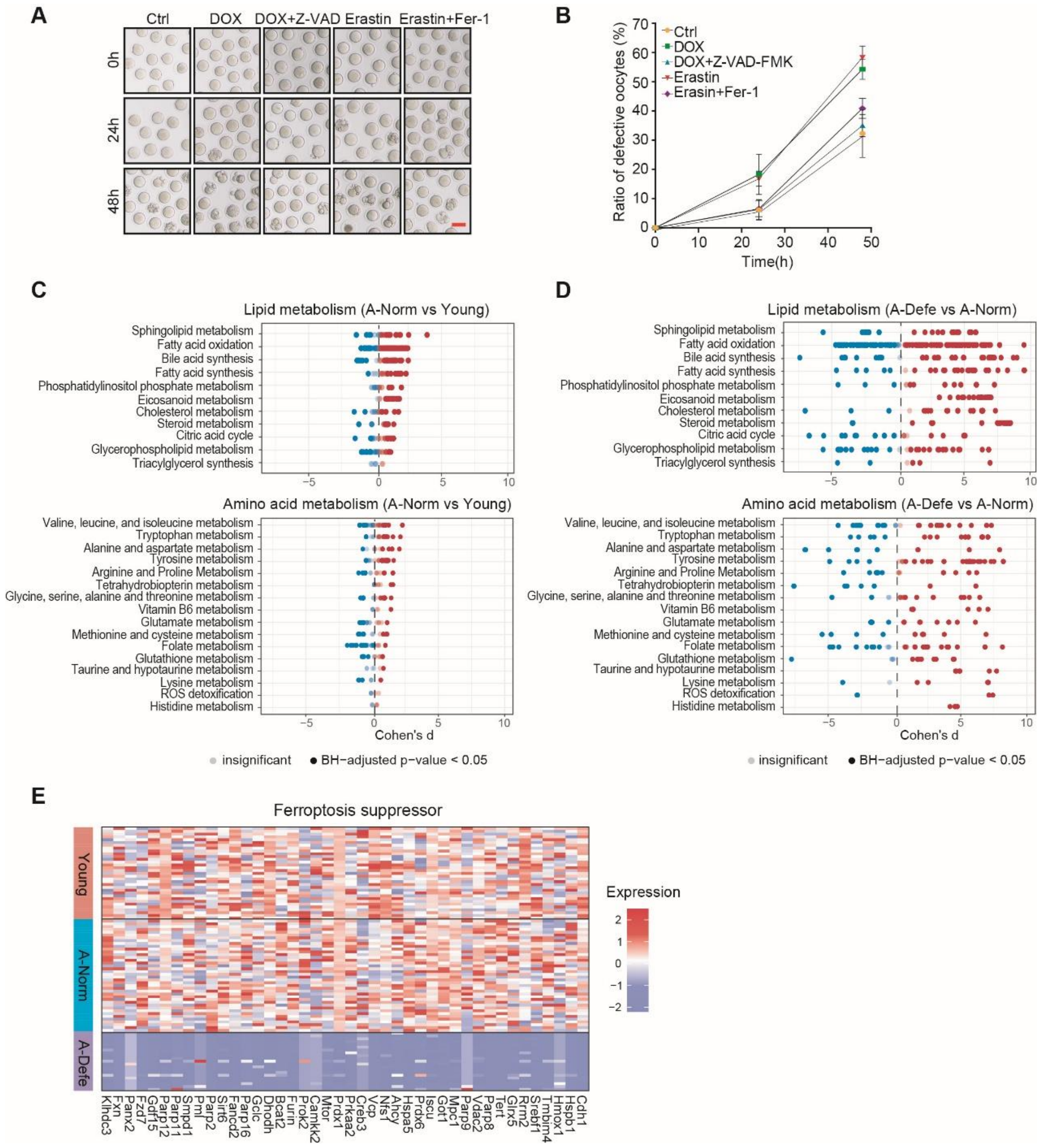


**Supplementary Fig. S3 | Both apoptosis and ferroptosis are involved in aged oocytes.** (**A**) Representative images showing the oocytes from young mice, treated DMSO (Ctrl), DOX, DOX with Z-VAD-FMK, Erastin and Erastin with Fer-1 for 0h, 24h, and 48 h. Scale bar, 100 μm. (**B**) Line graph comparing the defect rates of young oocytes following treatment with DMSO (n = 81), DOX (n = 93), DOX with Z-VAD-FMK (n = 107), Erastin (n = 114), and Erastin with Fer-1(n = 79) for 0 (H) 24 (H) and 48 h. Data are represented as the mean percentage (mean ± s.d.) from at least 3 independent experiments. (**C**) Dot plots showing Cohen's distance as a measure of dissimilarity between Compass scores of A-Norm and Young clusters, assigned to individual reactions (dots) within metabolic subsystems (y-axis)

in lipid metabolism (upper) and amino acid metabolism (lower). Metabolic reactions are subgrouped according to the Recon2 database and colored according to the Cohen's d statistic and adjusted p-value (depth of the color). (**D**) Analogue to (C) but comparisons between A-Defe and A-Norm clusters. Of note, the scale of the x-axes of (C, D) were kept the same. (**E**) Heatmap displaying the expression levels of selected ferroptosis suppressor genes. Oocytes were grouped according to the Young, A-Norm, and A-Defe clusters.

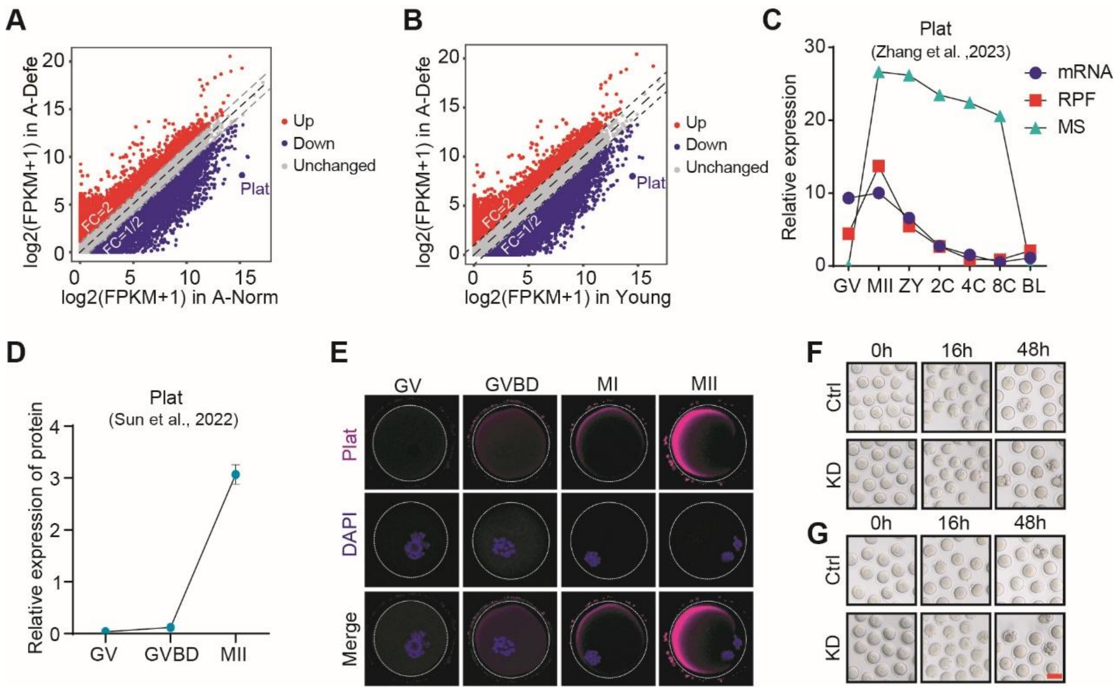


**Supplementary Fig. S4 | Elevated protein expression of *Plat* in MII-stage oocytes.** (**A**) Scatter plot comparing gene expression levels between A-Defe and A-Norm oocyte clusters. Highlighted in red or blue are genes exhibited increased or decreased expression over 2-fold in A-Defe oocytes. (**B**) Scatter plot comparing gene expression levels between A-Defe and Young oocyte clusters. Highlighted in red or blue are genes exhibited increased or decreased expression over 2-fold in A-Defe oocytes. (**C**) Line graph indicating the dynamics of Plat expression at the protein translation (based on ribosome protected fragments or RPF), and mRNA levels during oocyte maturation and early embryonic development. Protein expression was quantified using iBAQ, while RPF and mRNA levels were quantified using FPKM. All displayed expression values were log2 transformed. (**D**) Line graph illustrating the protein dynamics of tPA during oocyte maturation. (**E**) Representative images showing Plat expression and localization during oocyte maturation. Scale bar, 20 µm. (**F**) Representative images showing Plat-KD and Ctrl oocyte morphologies after Erastin treatment for 0 h, 16h, and 48 h. Scale bar, 100 µm. (**G**) Representative images showing Plat-KD and Ctrl oocyte morphologies after DOX treatment for 0h, 16h, and 48 h. Scale bar, 100 µm.

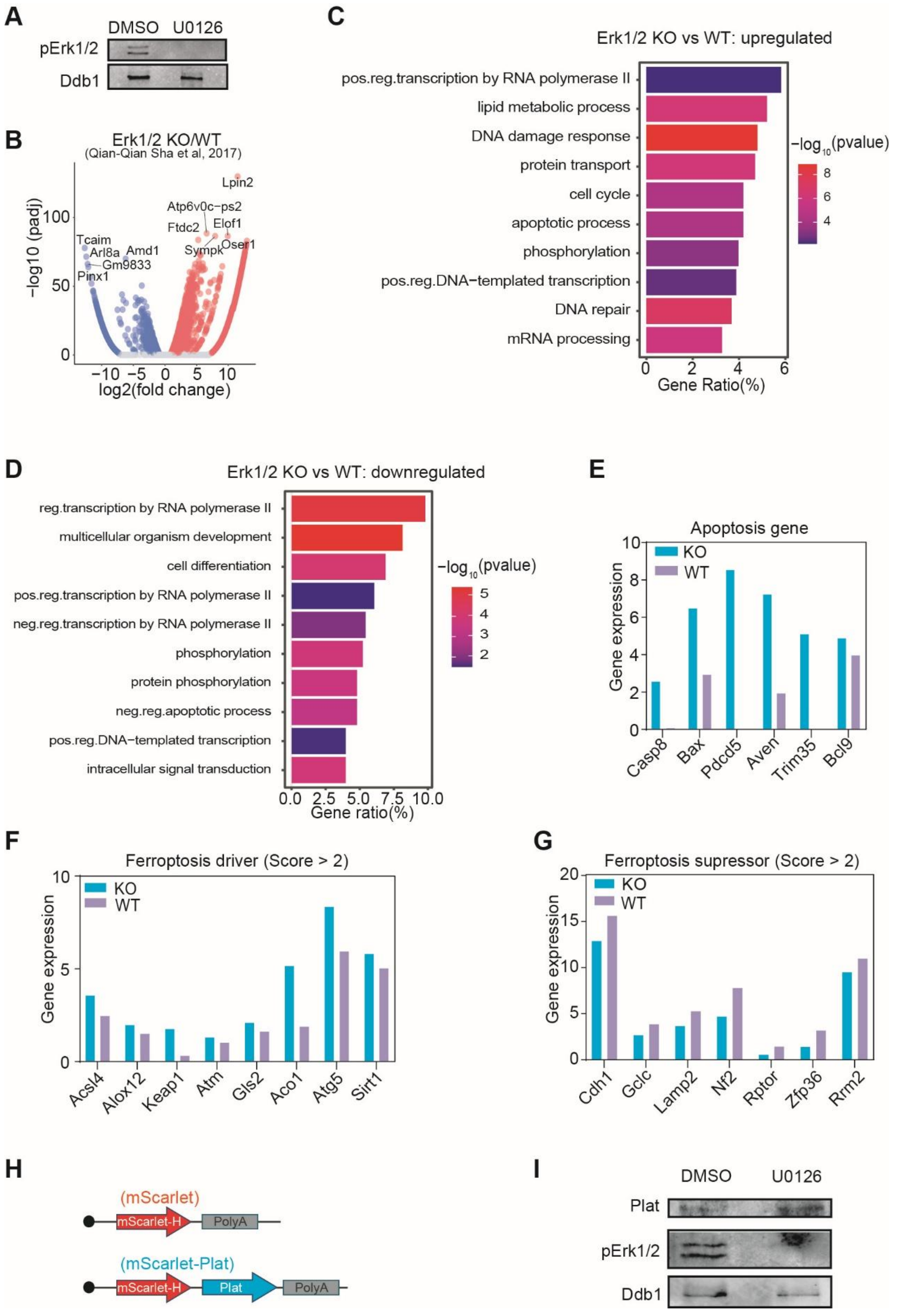


**Supplementary Fig. S5 | Plat is an upstream factor for Erk1/2 activation regulating oocyte death.** (**A**) Western blot analysis comparing pErk1/2 levels between oocytes treated with U0126 and DMSO. Ddb1 was used as a loading control. Total protein derived from 100 oocytes was loaded per lane. (**B**) Volcano plots showing differentially expressed genes in Erk1/2 knock-out (KO) oocytes compared to wild-type (WT) oocytes. Upregulated and downregulated genes (FC > 1.2 or FC < 1/1.2; and adjusted P value < 0.05) are depicted in red and blue, respectively. (**C**) Enriched GO terms in the upregulated genes in Erk1/2 KO oocytes compared to WT oocytes. (**D**) Enriched GO terms in the downregulated genes in Erk1/2 KO oocytes compared to WT oocytes. (**E**) Barplots comparing expression of apoptosis-related genes between Erk1/2 KO and WT oocytes. (**F**) Barplots comparing expression of ferroptosis drivers (score > 2 in FerrDb) between Erk1/2 KO and WT oocytes. (**G**) Barplots comparing expression of ferroptosis suppressors (score > 2 in FerrDb) between Erk1/2 KO and WT oocytes. (E-G) Gene expression levels are represented in transcripts per million (TPM). (**H**) Schematic diagram of in vitro transcribed mScarlet-Plat mRNA for microinjection. (**I**) Western blot analysis comparing Plat and pErk1/2 levels between oocytes treated with U0126 and DMSO. Ddb1 was used as a loading control. Total protein derived from 100 oocytes was loaded per lane.

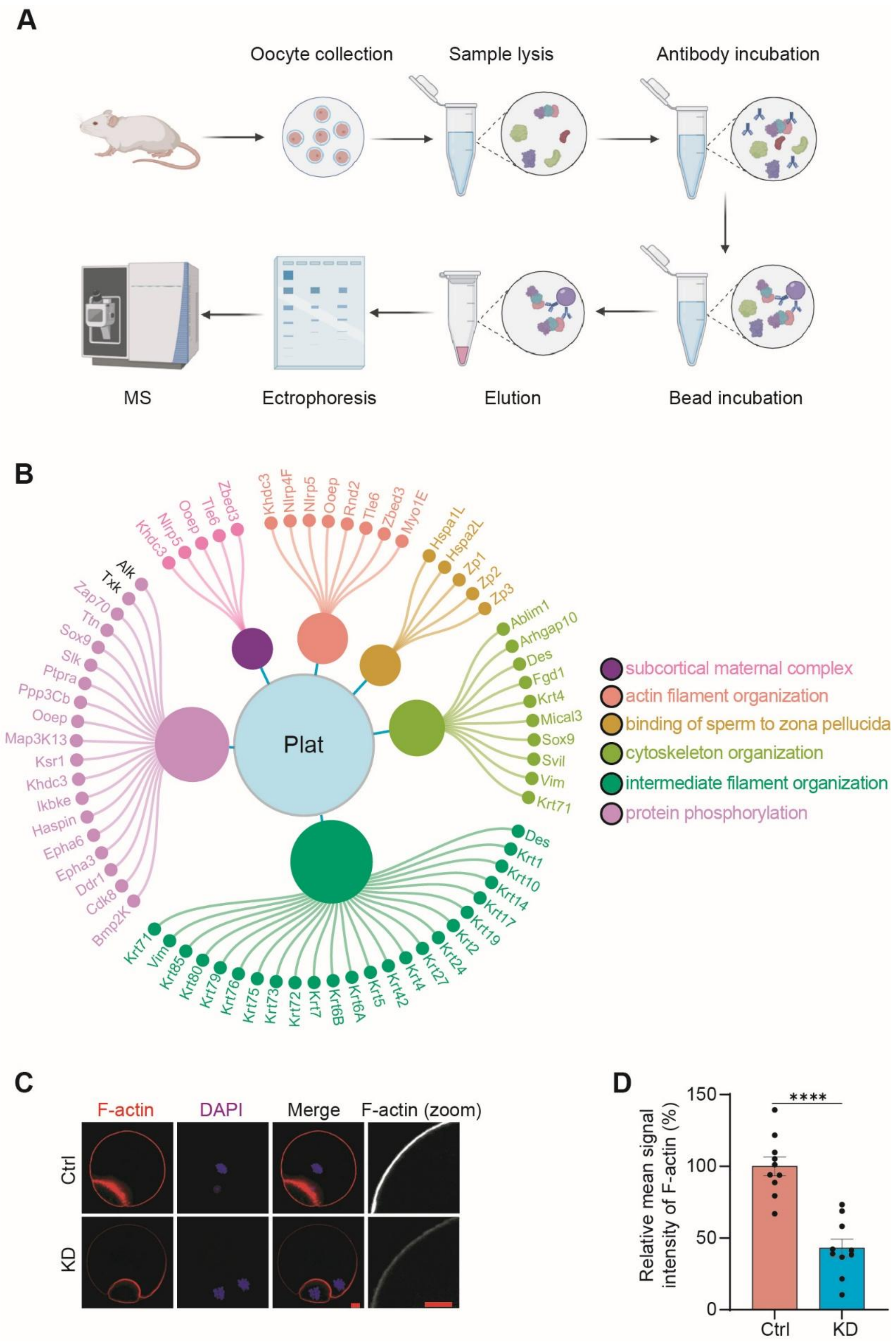


**Supplementary Fig. S6 | Protein interactome of tPA in oocytes.** (**A**) Workflow for tPA interatome identification by immunoprecipitation followed by mass spectrometry (IP-MS). Approximately 1500 mouse MII-stage oocytes were collected for immunoprecipitation (IP), followed by protein identification by mass spectrometry. (**B**) Network diagram showing the predominate categories of Plat-interacting proteins. Different colors depict distinct functional categories. (**C**) Immunofluorescence staining for F-actin in siCtrl (Ctrl) and Plat-knockdown (KD) oocytes (red), with DAPI-stained nuclei (blue). Magnified images of F-actin are displayed in gray. Scale bar, 10 μm. (**D**) Barplots comparing relative fluorescence intensity of F-actin between Ctrl (n = 10) and KD (n = 10) oocytes. Data are represented as the mean value (mean ± s.d.) of at least 3 independent experiments. The data are represented as the mean ± s.d. **** $P < 0.0001$ according to two-tailed unpaired t-test.
